# Tracking neural activity from the same cells during the entire adult life of mice

**DOI:** 10.1101/2021.10.29.466524

**Authors:** Siyuan Zhao, Xin Tang, Sebastian Partarrieu, Shiqi Guo, Ren Liu, Jaeyong Lee, Zuwan Lin, Jia Liu

## Abstract

Recording the activity of the same neurons over the adult life of an animal is important to neuroscience research and biomedical applications. Current implantable devices cannot provide stable recording on this time scale. Here, we introduce a method to precisely implant nanoelectronics with an open, unfolded mesh structure across multiple brain regions in the mouse. The open mesh structure forms a stable interwoven structure with the neural network, preventing probe drifting and showing no immune response and neuron loss during the yearlong implantation. Using the implanted nanoelectronics, we can track single-unit action potentials from the same neurons over the entire adult life of mice. Leveraging the stable recordings, we build machine learning algorithms that enable automated spike sorting, noise rejection, stability validation, and generate pseudotime analysis, revealing aging-associated evolution of the single-neuron activities.

## Main

Long-term stable recording^1–4^ of the same neuron at single-cell and single-spike resolution over the entire adult stage of life of behaving animals is important to understand how neural activity changes with learning and age^4,5,6,7^, to improve current brain-machine interface performance by reliably interpreting the brain’s behavioral and internal states^4,6^, and to study neurodegenerative diseases, aging-associated neurological disorders and cognitive decline^7,8^. Current implantable electronic and optical tools can record neural activity at single-cell and single-spike resolution but suffer from immune response and recording drift due to the mechanical and structural disparities between rigid electronic or optical devices and brain tissue^9,10^. Relative shear and repeat motion at the implanted interface keep changing the relative position between recording devices and recorded neurons. The proliferation of astrocytes and microglia form a ∼100 µm thick glial sheath that cause the death of neurons and isolate recording devices from neurons. Together, they lead to chronic instability of recordings. Optical imaging techniques are further limited by the light penetration depth and three-dimensional (3D) volumetric scanning across the 3D tissue due to optical aberration and attenuation^11^.

Miniaturized flexible electronics such as mesh nanoelectronics and thin-film probes have been utilized in *in vivo* electrophysiology^12–17^ given their unique mechanical properties. Mesh nanoelectronics provide a chronically stable, gliosis-free implantation over a few months through the incorporation of tissue-like structural and mechanical properties into nanoelectronics^18–20^. However, due to their mechanical flexibility, invasive methods such as syringe injection are required to implant the tissue-like electronics into the brain^18–20^. The relatively large mechanical damage from the implantation causes permanent damage to the neural network. In addition, implanted mesh nanoelectronics can only unfold in the cavities of the brain such as the subventricular zone, not in tissue-dense brain regions^18–21^. As a result, the bundled mesh nanoelectronics do not have the optimized mechanical flexibility required to interface with the brain tissue over long time periods^18–21^. On the other hand, flexible thin film brain probes lack the open mesh structures that allow for 3D integration with the neural network^22–24^. They also need to maintain mechanical strength to prevent damage of the probe during the implantation^22–25^, which can cause an immune response and probe drifting during long-term implantation. As a result, none of the existing technologies has demonstrated long-term stable tracking of the same neuron over the entire adult stage of life of a behaving animal.

Here, we solved this issue by implanting a fully unfolded tissue-like mesh nanoelectronics into the brain of a mouse. The fully unfolded mesh can form an interwoven structure within the neural network and eliminate the immune response and probe drifting, maintaining a long-term stable electrode-to-neuron interface at the single-cell level, thus enabling the same neuron to be recorded over the entire adult life of animals (Fig. 1a). To enable this method, we developed mesh nanoelectronics monolithically integrated with ultra-thin and releasable polymer shuttles through lithographic fabrication. We also incorporated unique polymer anchors and water-releasable structures, allowing for controllable, precise, and minimally invasive delivery of mesh nanoelectronics in the mouse brain. This implantation method can keep the designed open mesh structure of the device in the brain across multiple brain regions, including cell-dense regions. The open mesh allows the neural network to form seamlessly interwoven structures, provides a tissue-level flexible interface, and prevents the repeated micromotion and drift between recording electrodes and surrounding neural tissue during yearlong recording, thus allowing for a highly stable recording of neuron activities across multiple brain regions.

**Fig. 1.**
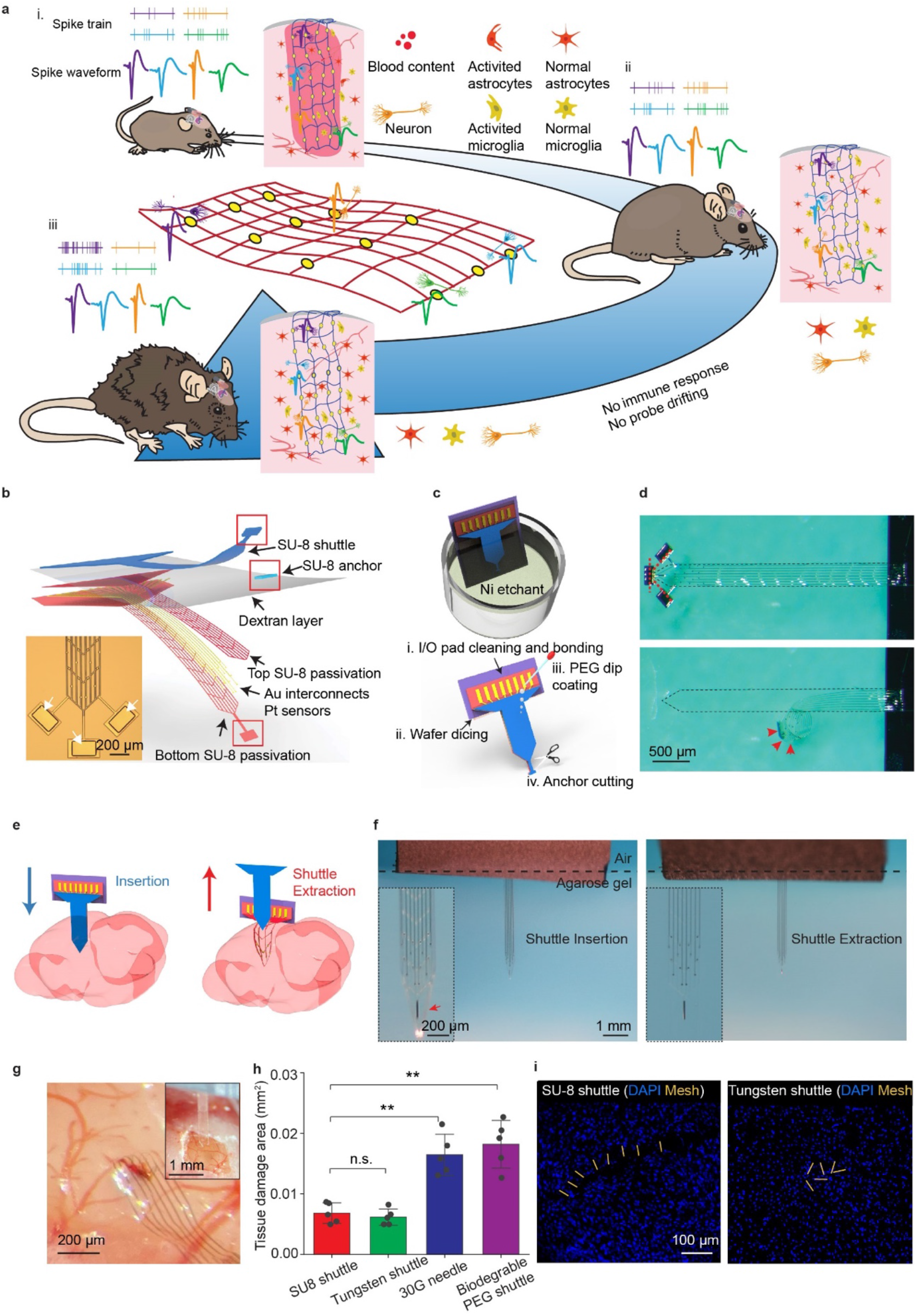
Minimally invasive implantation of tissue-level flexible mesh nanoelectronics in the brain. **a**, Schematics showing the long-term stable electrical recording of the same neuron over the entire adult life of mice enabled by the minimally invasively implanted and fully unfolded tissue-level flexible mesh nanoelectronics. Mesh nanoelectronics seamlessly integrate with neural networks without immune response. Single-cell electrophysiology from the same animal at (i) mature adult (3-6 months), (ii) middle (10-14 months), and (iii) old (18-24 months) stages are recorded. Colored waveforms represent different neurons that are stably recorded over the entire adult life of mice. **b**, Exploded view of the integrated mesh nanoelectronics showing the distinct material layers. The fully assembled nanoelectronics consisting of (from top to bottom) a 25-μm-thick polymer shuttle, a 20-μm-thick polymer anchor, a 3-μm-thick dextran dissolvable layer, a 450-nm-thick top SU-8 encapsulation layer, 50-nm-thick platinum electrodes and 70-nm-thick gold (Au) interconnects, and a 450-nm-thick bottom SU-8 encapsulation layer. Inset: bright-field (BF) microscopic image of mesh nanoelectronics connected with polymer shuttle through anchors (white arrows). **c**, Schematics showing stepwise releasing of mesh nanoelectronics from the substrate and shuttle. Mesh nanoelectronics was released from the fabrication substrate after removing the Ni sacrificial layer while connected with the polymer shuttle by the anchors (top). The released shuttle/nanoelectronics were cleaned for bonding and dicing, and then coated by the biodegradable PEG through dip-coating. After cutting the polymer anchors, the mesh nanoelectronics was released from the shuttle by dissolving PEG (bottom). **d**, Photographs showing the released polymer shuttle/mesh nanoelectronics hybrid from the substrate (top) and released mesh nanoelectronics from the shuttle (bottom). Red dashed lines and arrows highlight the cutting lines of the anchors and the released mesh nanoelectronics, respectively. **e**, Schematics showing the brain implantation process. **f**, *In vitro* images of mesh nanoelectronics implantation in a 0.6% agarose gel. Insets: zoom-in images showing the released mesh nanoelectronics maintain the unfolded structure and implantation location after withdrawing the polymer shuttle. **g**, Photograph showing the representative brain implantation with minimal tissue damage. Inset: ultrathin polymer shuttle-enabled implantation. **h**, Statistical analysis of tissue acute damage zone of different implantation methods. Data represented as mean ± SD, individual data points are overlaid (*\*\*p* < 0.01, two-tailed unpaired *t* test, *n* = 5). **i**, Representative images of 20-μm-thick horizontal brain slices showing the acute mechanical injuries by the ultrathin-polymer shuttle (left) and 50-μm diameter tungsten shuttle (right). Yellow and blue represent mesh nanoelectronics and DAPI, respectively.

By optimizing the size of the implanted nanoelectronics, we achieved stable tracking of the same neuron over the entire adult life of mice until their natural death (*i.e.,* 5-18 months for mouse #1, 5-20 months for mouse #2, and 5-19 months for mouse #3), confirmed by vigorous statistical tests^3,18–21,26^. Tracking the whisker-stimulation-evoked single spikes from the barrel cortex indicated that the electrode embedded in a single whisker barrel does not drift over the animal’s adult life. Using the first several months’ recording data to train an autoencoder^27^, a machine learning (ML) tool for representation learning, we further confirmed the stable recording, which also allowed for fully automatic spike sorting, noise rejection, and stability analysis over the adult life of the mice. Finally, ML-based pseudotime analysis of single-unit waveforms identified several neurons with age-dependent changes in electrical activities.

## Results

### Monolithically integrated mesh nanoelectronics and ultra-thin shuttle

To implant completely open and unfolded mesh nanoelectronics into the brain, we integrated the mesh nanoelectronics monolithically with a releasable, ultra-thin polymeric shuttle using standard photolithography procedures (Fig. 1b, Extended Data Fig. 1, 2a-k, Methods). The mesh nanoelectronics were fabricated as described in previous reports^18–20^. Briefly, 16 or 32 15-μm-diameter electrodes were connected by SU-8 encapsulated Cr/Au interconnects to Cr/Au input/output (I/O) pads. The encapsulated interconnects were 10-μm wide and <1-μm thick, forming a mesh network with a 2D filling ratio at 73.3%, which yielded an effective bending stiffness of 1.26×10^−15^ N·m^2^. The mesh nanoelectronics were partially fabricated on the top of a Ni sacrificial layer. Next, a 25-μm-thick polymer shuttle was defined on the top of the mesh nanoelectronics with a 3-μm-thick water-soluble dextran and 20-μm-thick polymer anchors. These polymer anchors (Fig. 1b, inset) connected the mesh nanoelectronics with the polymer shuttle through the dextran layer (Fig. 1b, red box). After the integrated device was released from the substrate (Fig. 1c, top) by removing the Ni sacrificial layer, the anchor kept the pattern of the mesh nanoelectronics on the polymer shuttle. Then, a few drops of 10 wt% PEG (35 kDa) was coated to reinforce the bonding between the mesh nanoelectronics and polymer shuttle (Fig. 1c, bottom), as well as enhanced the temporary stiffness and provided protection to the mesh nanoelectronics during implantation. The biodegradable PEG adhesion layer has a sub-micron thickness; thus, the surgical footprint is mostly affected by the thin polymer shuttle. After removing the anchor connection (Fig. 1c, bottom), the mesh nanoelectronics can be readily released from the polymer shuttle by dissolving the PEG in an aqueous solution (Fig. 1d).

The polymer shuttle was used to guide the implantation of the mesh nanoelectronics into the brain tissue (Fig. 1e, left). After the integrated brain probe reached the targeted position, saline was applied to quickly dissolve the PEG and to release the mesh nanoelectronics from the polymer shuttle, which was subsequently withdrawn from the brain tissue (Fig. 1e, right). We characterized the implantation procedure in a transparent 0.6 wt% agarose gel-based brain phantom with mechanical properties comparable to that of the brain tissue^28^. At a typical implantation speed of 100 μm/s, we did not observe any buckles on the probes (Fig. 1f, left). After insertion, 1× phosphate buffered saline (PBS) solution was applied to dissolve the PEG/dextran adhesive layer. The shuttle was then withdrawn at a speed of 10 μm/s. After withdrawing the shuttle, the mesh nanoelectronics still maintained its implantation location and open mesh structure without any deformations (Fig. 1f, right). We tested the yield of the implantation in brain phantoms with various speeds. We achieved a 93.3% yield for insertion and extraction of 16-channel mesh nanoelectronics at 100 μm/s (Extended Data Fig. 2l, m). There was no significant change of implantation yield with different size mesh electronics of 32 channels (*n* = 3, *p* > 0.05, Extended Data Fig. 2l, m).

The optimized implantation procedure was then used for mouse brain implantation. Figure 1g shows typical implantation of 300-μm-wide 16-channel mesh nanoelectronics in the anesthetized mouse brain (Methods). The probe (Fig. 1g, inset) can be easily implanted with the polymer shuttle withdrawn by the same conditions tested for the phantom gel. Damages to the blood vessels were minimal throughout the imaging-guided implantation. To evaluate acute tissue damage, cell loss, and mesh nanoelectronics distribution, we imaged the *post hoc* fixed and stained tissue slices immediately after implantation (Fig. 1h). Acute damaged area with the thin-shuttle was approximately 0.0068 ± 0.0016 mm^2^ (mean ± SD, *n* = 5, Fig. 1h), which is significantly smaller than those from previous reported implantations using syringe-injection^18,19^ (0.0164 ± 0.0033 mm^2^, mean ± SD, *p* < 0.01, *n* = 5, Fig. 1h) and biodegradable shuttles^21,29^ (0.0182 ± 0.0039 mm^2^, mean ± SD, *p* < 0.01, *n* = 5, Fig. 1h). While the tissue damage showed no significant difference compared with samples implanted by 50-μm diameter tungsten wire^25^ (0.0062 ± 0.0014 mm^2^, mean ± SD, *n* = 5, Fig. 1h), the cross-sectional images of brain slices with implants showed the clear unfolded mesh structures vs. bundled ribbons by tungsten probe-based delivery (Fig. 1i). Compared with a previously demonstrated implantation method for flexible neural probes^18–21,25,29,30^, the integrated 25-μm-thick polymer shuttle drastically reduces tissue displacement during implantation as well as maintains the designed open structure with nearly 90% implantation yield. Moreover, the presented method involves minimal manual manipulations since the ultra-flexible nanoelectronics was pre-attached to the thin-shuttle with the lithography process. On average, it took less than 3 min to assemble one mesh nanoelectronics (more than 20 nanoelectronics per hour), including sub-micro-thick PEG coating, anchor dicing, and additional packaging with a success rate approaching 100%.

### Unfolded mesh nanoelectronics structure 3D interwoven with the neural network

We implanted mesh nanoelectronics with different sizes across multiple brain regions. Each mesh structure has an ultra-small cross-section of 10×1 μm^2^. The longitudinal bending stiffness of each individual mesh structure reached 1.26×10^−15^ N·m^2^, which is comparable to that of brain tissue and orders of magnitude lower than *state-of-the-art* probes (*i.e.,* ultrasmall carbon^31–33^, polyimide^34^ and elastomer-based ‘e-dura’ probes^35^). Rhodamine 6G was added to the SU-8 encapsulation layer, enabling the imaging of the mesh structure in the brain. To explore the potential capability of the implantation, Figure 2a shows a 2-mm-wide, 3-mm-long mesh nanoelectronics implanted into a mouse brain across cortex, hippocampus, and thalamus regions. The size of this device can potentially include 1,024 recording sites through 3D stacking of electrodes^36^ (Fig. 2a-d, Methods). A representative 3D reconstructed image of the mesh nanoelectronics in the brain tissue at 6-week post-implantation (Fig. 2a) showed the fully unfolded, open mesh structure interweaving with neurons and astrocytes across multiple brain regions (*i.e.,* cortex, hippocampus, thalamus, etc.). A slight bending of the mesh suggested that the tissue-like nanoelectronics were flexible within the tissue. A zoomed-in view of the hippocampus CA1 region (Fig. 2b) shows a smooth distribution of neurons and astrocytes across the mesh. Notably, neurons in the cell-dense region (hippocampus) can still penetrate the open mesh structure (Fig. 2c), forming an intertwined tissue-nanoelectronics interface. Figure 2d shows that the size of the recording electrode (white dashed circles) is comparable to the size of the soma. The subcellular feature size, tissue-level flexibility, and 3D interwoven network collectively eliminated the micromotion between the functional electrode and recorded neurons^37^. Additional replications of mesh nanoelectronics with different sizes were implanted in mouse brains and are shown in Extended Data Fig. 3.

**Fig. 2.**
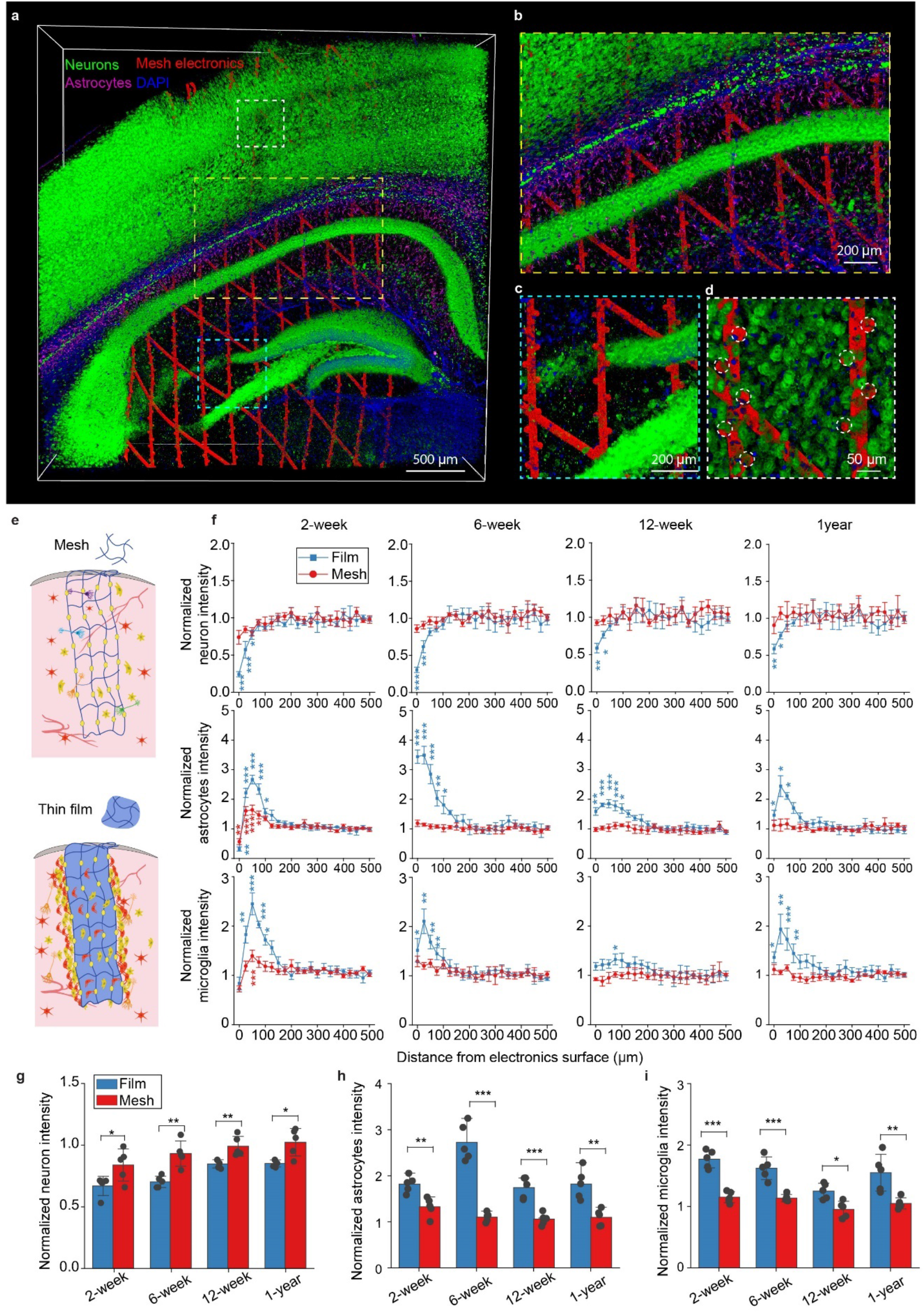
Unfolded mesh nanoelectronics seamlessly integrating with the neuron network across multiple brain regions. **a**, Representative 3D reconstructed confocal fluorescence imaging of 600-μm-thick brain tissue implanted with 2-mm wide mesh nanoelectronics for 6 weeks. Green, purple, blue, and red label neurons, astrocytes, nuclei, and mesh nanoelectronics. **b**-**d**, Zoom-in views of the regions highlighted by white (**b**), cyan (**c**), and yellow (**d**) dashed boxes in (**a**), showing the seamless integration of the mesh with the neural network. Neuron-like electrodes are highlighted by white dashed circles in (**d**). **e**, Schematics illustrating 1-μm-thick mesh (top) and thin-film (bottom) nanoelectronics implanted and unfolded inside brain tissue for the long-term immune response characterization. **f**, Normalized fluorescence intensity as a function of distance from the mesh/thin-film electronic and tissue boundary at 2 weeks, 6 weeks, 12 weeks, and 1-year post-implantation. The relative signal was obtained by normalizing the fluorescence intensity with the baseline value defined as the fluorescence intensity over a range of 525–550 μm away from the electronics. Data represented as mean ± SEM (intensity compared with that of distance at 500-μm, *\*p* < 0.05, *\*\*p* < 0.01, *\*\*\*p* < 0.001, two-tailed unpaired *t* test, *n* = 5). **g-i**, Normalized neuron (**g**), astrocytes (**h**), and microglia (**i**) intensity and neuron density within 100-μm from the electronic surface. Data represented as mean ± SD, individual data points are overlaid (*\*p* < 0.05, *\*\*p* < 0.01, *\*\*\*p* < 0.001, two-tailed unpaired *t* test, *n* = 5).

Next, we performed longitudinal immunostaining characterizations to assess the distribution of key cell types around mesh nanoelectronics over the time course of implantation. To demonstrate that the open mesh structure reduces immune responses during chronic implantation (Fig. 2e, top), thin-film nanoelectronics with the same dimensions as mesh nanoelectronics were used (Fig. 2e, bottom) as control and contralaterally implanted in the same mouse brain. The bending stiffness of the thin film nanoelectronics is only slightly higher than that of the mesh nanoelectronics (39.8× 10^−15^ N·m^2^ vs. 1.26×10^−15^ N·m^2^, Methods). The brain tissue was harvested and sliced for immunostaining 2-, 6-and 12-week, and 1-year post-implantation. Horizontal slices were stained with cell-type-specific protein markers for imaging of neurons, astrocytes, and microglia (Extended Data Fig. 4). We quantitatively analyzed horizontal brain slices implanted with 16-channel, 300-μm-wide film/mesh nanoelectronics (Fig. 2f-i, Methods). Protein marker signals were calculated by normalizing the fluorescence intensity around the implantation site with the baseline value defined as the average fluorescence intensity over a range of 525-550 μm away from the nanoelectronics. Statistical analysis demonstrated a significant degradation of neuron density (NeuN) and an enhancement of astrocytes and microglia intensity (GFAP and Iba-1, respectively) near the thin-film nanoelectronics at all time points (*p* < 0.05, *n* = 5, Fig. 2f). These results proved that the thin-film nanoelectronics can still trigger the proliferation of astrocytes/microglia and reduced the neuron density at the nanoelectronics-brain interface. Meanwhile, the open mesh nanoelectronics introduced minimal damage to the surrounding neurons and negligible immune response. Importantly, the result demonstrated that no significant neuron loss was detected at 2-week post-implantation for mesh samples (Fig. 2f), suggesting minimal acute damage from the thin-polymer shuttle. In addition, the neuron density near the mesh surface remained the same at one-year post-implantation (Fig. 2f).

We further calculated the normalized intensity of neural cell fluorescence signals within regions 100-μm away from the nanoelectronics to assess neuron loss and inflammation reaction at the different post-implantation periods (Fig. 2g-i). The mesh nanoelectronics samples showed a neuron density of 83.9 ± 13.0% (mean ± SD, *n* = 5, Fig. 2g) at 2 weeks, which is greater (*p* < 0.001, *n* = 5, Fig. 2g) than that from the thin film nanoelectronics (66.9 ± 7.8%, mean ± SD, Fig. 2g) in the contralateral brain slices. Compared to the non-implanted regions, neuron intensity of mesh nanoelectronics samples increased to 93.1 ± 10.2%, 99.1 ± 8.0%, and 102.4 ± 11.2% (mean ± SD, *n* = 5, Fig. 2g) 6 weeks, 12 weeks, and 1-year after implantation, respectively. On the contrary, thin-film nanoelectronics samples showed significant neuron loss for the same periods (70.2 ± 4.6%, 84.8 ± 3.3%, and 85.2 ± 2.3% at 6 weeks, 12 weeks, and 1-year after implantation, respectively. *p* < 0.05, *n* = 5, Fig. 2g). The intensity of astrocytes and microglia around the mesh slightly increased at 2 weeks (115.1 ± 9.1% at microglia, 132.6 ± 20.5% at astrocytes, mean ± SD, *n* = 5, Fig. 2h, i) and then reduced at 6 weeks (113.6 ± 5.8% at microglia, 110.1 ± 10.0% at astrocytes, mean ± SD, *n* = 5, Fig. 2h, i). Moreover, continuous monitoring of the inflammation around the mesh nanoelectronics revealed nearly normal immune cell distribution at 12 weeks (95.3 ± 13.1% at microglia, 105.5 ± 12.7% at astrocytes, mean ± SD, *n* = 5, Fig. 2h, i), and even up to one year (104.9 ± 9.0% at microglia, 109.5 ± 17.9% at astrocytes, mean ± SD, *n* = 5, Fig. 2h, i). We attribute the little-to-no immune response of the mesh nanoelectronics to the ultra-flexible open structure that is imperceptive to surrounding brain tissue, neurons, and the cells involved in inflammation. Compared with the open mesh structure, the thin-film nanoelectronics implantation introduced significantly higher levels of astrocytes (181.9 ± 17.9%, 272.6 ± 40.3%, 174.4 ± 22.8% and 182.1 ± 32.7% at 2-, 6-, 12-and 1-year post-implantation, respectively, mean ± SD, *n* = 5, Fig. 2h, i) and microglia aggregation (177.2 ± 14.5%, 162.3 ± 18.4%, and 125.1 ± 12.6% and 154.9 ± 30.0% at 2-, 6-, 12-and 1-year post-implantation, respectively. Mean ± SD, *n* = 5, Fig. 2h, i) over the same period (*p* < 0.05, significant enhancement compared with the open mesh at all time points, Fig. 2h, i). Notably, we can still observe the proliferation of astrocytes and microglia around the thin-film nanoelectronics at one year post-implantation. Together, these results demonstrate that open mesh nanoelectronics introduce little-to-no inflammation and mechanical damage to the surrounding tissues as compared with thin-film nanoelectronics over yearlong implantation.

### Long-term stable recording at single-cell resolution across multiple brain regions

To test the stability of the recording, we implanted 600-μm-wide mesh nanoelectronics with 32 channels and 300-μm-wide mesh nanoelectronics with 16 channels into multiple mouse brain regions for head-fixed behaving electrophysiology (Methods). Electrodes were implanted into different brain regions including the somatosensory cortex and striatum (32-channel, mesh nanoelectronics#1, Fig. 3a); red nucleus, interstitial nucleus, and ventral tegmental area in midbrain (32-channel mesh nanoelectronics#2, Fig. 3a); and visual cortex and hippocampus (16-channel mesh nanoelectronics#3, Fig. 3a). Putative individual neurons were isolated using *Waveclus3*^38^ (Methods). Intrinsic spike waveform variability from the superficial (mesh nanoelectronics #1, #3, Fig. 3b, d) and deep (mesh nanoelectronics #2, Fig. 3c) brain regions is consistent with different putative neuron types in each brain region^39,40^. Moreover, the hippocampus recordings show higher neuron yield per electrode (2.0 ± 0.4 neurons, median ± 1.5 interquartile range, *n* = 8, Fig. 3e), spike amplitude (148.146 ± 77.4 μV, median ± 1.5 interquartile range, *n* = 8, Fig. 3f) and firing rate (17.3 ± 7.6 spike/s, median ± 1.5 interquartile range, *n* = 8, Fig. 3g) as compared to other recorded brain regions (primary somatosensory cortex, striatum, midbrain, primary visual cortex, Fig. 3e-g).

**Fig. 3.**
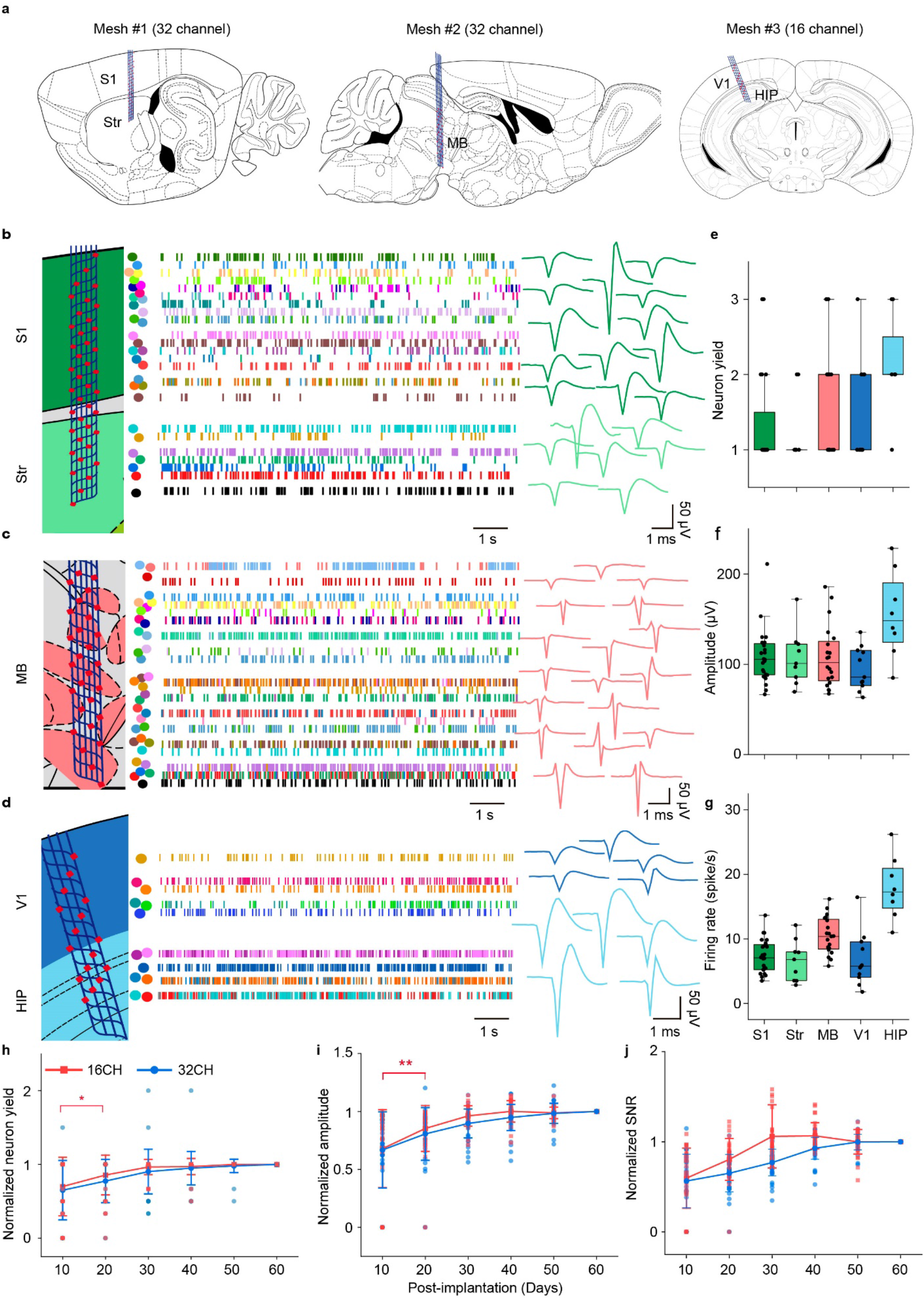
Chronically stable recording across multiple brain regions. **a**, Schematics illustrating the representative brain regions implanted with mesh nanoelectronics for chronic recording. S1, primary somatosensory cortex; Str, striatum; MB, midbrain; V1, primary visual cortex; HIP, hippocampus. **b**-**d**, Approximate mesh locations overlaid on the Allen Mouse Brain Atlas (left) and representative spiking raster (middle) and single-unit waveform (right) from two 32-channel (**b**, **c**) and one 16-channel (**d**) mesh nanoelectronics in head-fixed behaving animals at 60 days post-implantation. Colored dots and blocks indicate individual neurons and spike times, respectively (middle). Representative average single-unit action potential waveforms were extracted from a 2-min recording session (right). **e**-**g**, Quantification of sorted neuron yield per electrode (**e**), waveforms amplitude (**f**), and firing rate (**g**) across 5 brain regions in a 2 min recording session. Box plots show median and quartile range (whiskers denote 1.5× the interquartile range). Individual data points are overlaid (*n* = 29 electrodes from three 16-channel mesh nanoelectronics and *n* = 43 electrodes from two 32-channel mesh). **h-j**, Normalized sorted neuron yield (**h**), amplitude (**i**), and signal-to-noise ratio (SNR) (**j**) over the time course of 60 days. Data represented mean ± SD, individual data points are overlaid (*n* = 29 electrodes from three 16-channel mesh nanoelectronics and *n* = 43 electrodes from two 32-channel mesh nanoelectronics. *\*p* < 0.05, *\*\*p* < 0.01, comparison of different days within 16-,32-channel mesh electronics, two-tailed unpaired *t* test).

Next, we evaluated the long-term stability of recordings from 32-channel and 16-channel mesh nanoelectronics from 5 independent animals (*n* = 43 electrodes from two 32-channel and *n* = 29 electrodes from three 16-channel mesh nanoelectronics). 72 putative individual neurons from multiple regions were recorded 10 days post-implantation, which increased to 115 putative individual neurons after 60 days. Both 16-channel and 32-channel mesh nanoelectronics show low noise level and high signal to noise ratio (SNR) at 60 days post-implantation in behaving animals (16-channel: 9.97 ± 1.72 µV at noise level, 12.67 ± 6.34 at SNR, *n* = 29 electrodes; 32-channel: 9.27 ± 2.10 µV at noise level, 13.25 ± 5.94 at SNR, *n* = 43 electrodes, mean ± SD). The statistical results (Fig. 3h-j) showed that the normalized neuron count per electrode, average amplitude, and SNR of 300-µm-wide,16-channel mesh nanoelectronics increased over the first 30 days of implantation and then stabilized (*n* = 29 electrodes from three 16-channel mesh nanoelectronics). These parameters from the 600-µm-wide, 32-channel mesh nanoelectronics stabilized at 50-day post-implantation, suggesting the potential device size-related effect on the signal stability (*n* = 43 electrodes from two 32-channel mesh nanoelectronics). These results contrast with reports from previous brain probes where amplitudes, SNR, and neuron counts degrade weeks after implantation^3,4,37^, suggesting that the unfolded, open mesh nanoelectronics formed a long-term stable interface with neurons and tissue.

### Tracking the same neuron’s activity over the entire adult life of mice

Two mice implanted with 16-channel mesh nanoelectronics and one mouse with 32-channel mesh nanoelectronics were recorded monthly until their natural death (5-18 months for mouse #1, 5-20 months for mouse #2, and 5-19 months for mouse #3). 82.8 ± 6.2 % neurons are stably recorded (mean ± SD, compared to the first recording session). Compared to mice of 5 months with glossy brown fur, the aged mice of 18-months with mesh electronics implanted exhibited normal and healthy aging, including weight gain, barbering around the eyes, and thinning and grey fur in the dorsal back skin^41^ (Extended Data Fig. 5a-e). Statistical analysis revealed that there was a significant increase of gray hairs and a decrease of black hairs in aged mice when compared to the mature adult mice (*p* < 0.05, *n* = 3, Extended Data Fig. 5f). The electrode interfacial impedances exhibited relatively constant values of 920.2 ± 107.2 kΩ vs. 857.2 ± 85.7 kΩ at months 6 vs.18 (mean ± SD, *n* = 30, Extended Data Fig. 6a), indicating stable electrical and mechanical properties of the mesh nanoelectronics^42^.

We first assessed the stability of the signals by spike sorting and statistical analyses. Spike waveforms were projected to a 2D embedding space for stability validation by using UMAP-^26^ and PCA-based dimension reduction algorithms. PCA is commonly used to define the number and stability of recorded single-neuron signals over time^19,20,31^. UMAP is a non-linear ML-based dimension reduction algorithm that can learn a low-dimensional embedding space to preserve as much of the local and more of the global data structure than linear dimension reduction algorithms such as PCA. 26 neurons were isolated across all recording sessions starting from 5 months and lasting until natural death. The clusters for each sorted spike in both UMAP and PCA embeddings show nearly constant positions and well separated from each other in the first and second component plane (UMAP1-UMAP2 and PC1-PC2) through >1-year recordings (Fig. 4a, Extended Data Fig. 6b, Methods). In addition, the corresponding single-unit waveforms’ shapes (Fig. 4b), as well as their firing dynamics (*i.e.*, interspike interval) were stable (Extended Data Fig. 6c, Methods). Auto-correlation analysis showed that single-unit waveforms were highly similar and almost indistinguishable to themselves (0.90 ± 0.14 across all recording sessions from 3 mice, mean ± SD, Fig 4c, Methods). L-ratio^43^ and silhouette score^44^ analysis (Fig. 4d, Methods) confirmed good unit separation and accurate identification of individual neurons, demonstrating that the signals were sufficiently separated to permit isolation of single units. Statistical analysis on single-unit recording stability examined by five waveform features (amplitude, duration, peak-trough ratio, repolarization slope, and recovery slope)^40^ and signal-to-noise ratio (SNR) showed that their average values were nearly constant and the majority (79% from 3 mice) of neuron waveform features did not change significantly over time (*p* > 0.05 two-sided *t* test, Fig. 4e-h, Methods), demonstrating that the neuron spikes showed consistent characteristic features over time. Notably, the consistent signal-to-noise ratio (SNR) demonstrated that the electrode-to-cell interface was not degraded during the entire period (Fig. 4f). Collectively, all these results indicate that the waveforms were stably recorded from the same neuron over the entire recording period.

**Fig. 4.**
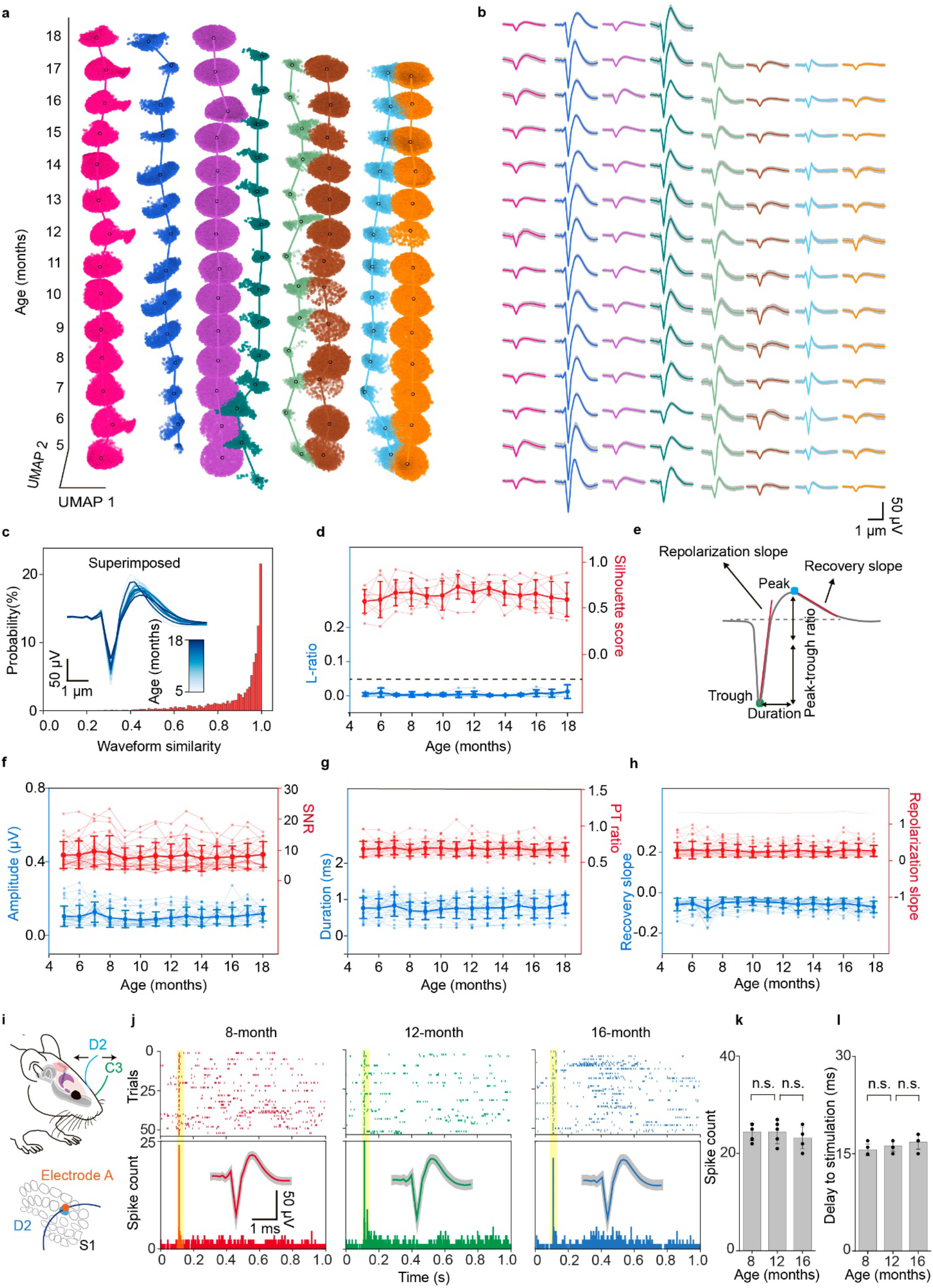
Stably tracking the single-unit action potential from the same group of neurons over the entire adult life of mice. **a**, Time evolution of single-units clustered by *Leiden* over the entire adult life of mice from the mature adult (5 months) to the aged (18 months) stage. The x- and y-axes denote the first and second uniform manifold approximation and projection (UMAP) dimensions, respectively, and the z-axis denotes mouse age in months. Color scheme maintained throughout. **b**, Time course analysis of the average waveforms of single-unit action potential in the *Leiden* clustering results in (**a**). Waveform represented mean ± SD. **c**, Waveform correlations for every single-unit across all recording sessions (*n* = 26 units from 3 mice). Inset: Overlaid average waveforms across all recording session for the representative putative same neuron. **d**, Likelihood-ratio (L-ratio) and silhouette score showing the clustering quality for the single unit action potentials from identified neurons. Data represented mean ± SD, individual data points are overlaid. The constant dashed line is 0.05 L-ratio, commonly taken as a threshold of high cluster quality. **e**, Schematic showing features extracted from the single-unit action potential waveform used for the analysis in (**f**-**h**). **f**-**h**, mean ± SD (individual data points are overlaid) of features illustrated in (**e**) and signal-to-noise ratio (SNR) over three mice life recording. In order, amplitude, SNR, duration, peak-trough ratio (PT ratio), recovery slope, and repolarization slope are shown. **i**-**l**, Long-term stable recording of the behavior-associated neuron. i, Schematic diagram of whisker deflection. An individual vibrissa was deflected in the rostral-caudal plane using a computer-controlled galvanometer system. D2 and C3 whiskers are labeled in blue and green, respectively (top). Schematic diagram of whisker barrel arrangement in S1 (bottom). Evoked spike from electrode A was associated with D2 whisker deflection. Control experiments were shown in Extended Data Fig. 7. **j**, Raster plot and peri-stimulus time histogram (PSTH, 1 ms bin size) of the single unit identified from electrode A response to D2 whisker deflection from 8- to 16-month recording. Inset: average single-unit waveforms from recordings in the S1 in response to D2 whisker deflection over time, Waveform represented mean ± SD. **k**, Population data showing the number of spikes (out of 55 trials) evoked by whisker deflection over the time course of implantation. **l**, Time delay of the evoked spikes to the stimulation over the time course of implantation (n.s.: not significant, two-tailed unpaired *t* test, *n* = 5). Individual data points are overlaid.

In addition to recording of spontaneous activity, we validated the stability of recording by examining the stable recording of whisker stimulation-elicited neuron activities^32,45^. Specifically, we identified one electrode (electrode A) on the mesh nanoelectronics as being close to a D2 barrel neuron in primary somatosensory cortex (S1 cortex) by successful recording of the whisker-stimulation-elicited single-unit spikes (Fig. 4i-l). Recording from stimulation of other whiskers (*e.g.*, C3) or another electrode (*e.g.*, electrode B that is close to electrode A) away from the D2 barrel field were used as control (Extended Data Fig. 7). We performed the whisker deflection with a galvanometer-driven stimulation contralateral to the implant (Fig. 4i). A 1 Hz, 900 deg/s deflection was applied to the targeted whisker (Methods). The raster plot and peri-stimulus time histogram (PSTH) of this single-unit recording showed that the electrode A of the mesh nanoelectronics can record strong and rapid neuron firing in response to the principal whisker D2 deflection (Fig. 4j). We can record well defined neuron activity and waveforms from 8 months to 16 months (Fig. 4j). Spikes observed from the electrode A with C3 whisker deflection (Extended Data Fig. 7a-c) or from electrode B with D2 whisker stimulation (Extended Data Fig. 7d-e) showed no correlation with the whisker stimulation. Notably, Figure 4k and l showed that the evoked spike count and the time delay to the stimulation exhibited no significant change over time (*p* > 0.05, *n* = 5). These behavior-triggered electrical activities further demonstrated the capability of this method to track the same neuron during the adult lifetime of mice.

### ML-based validation and analysis

We further applied an unbiased, autoencoder-based^27^ automated neuronal signal processing and analysis to benchmark the stability of the signal. Autoencoders is a self-supervised learning algorithm, which is only able to meaningfully reconstruct data similar to what they have been trained on, thus providing an unbiased way to examine the stability of the recording. We trained three two-headed autoencoders by using detected spike waveforms, corresponding electrode information and sorted neuron labels from the first 6-month recording data of the three mice. Each autoencoder i) learned nonlinear dimension reduction transformations compressing spike waveforms into a 2D embedding space, ii) classified the spike as a specific neuron in the training data, and iii) reconstructed the embeddings back to the input data space (Fig. 5a). The classification and reconstruction were simultaneously optimized during the training of the autoencoder, enabling the autoencoder the capabilities of spike sorting, postprocessing, and stability verification (Extended Data Fig. 8a, Methods) at the same time. Notably, the autoencoder showed much faster classification speed and higher reconstruction accuracy compared with UMAP and random forest^46^ classifier-based spike sorting (Extended Data Fig. 8b, Methods). While we used the autoencoder trained with the first six-month recording to achieve the best performance, we found that a two-month recording dataset is sufficient to train the model with only about 4% accuracy decrease (Extended Data Fig. 8c). The waveforms of the remaining 8-month recording data can still be classified and reconstructed (Fig. 5b and Extended Data Fig. 8d) with the classification accuracy (Fig. 5c, Extended Data Fig. 8e) reaching 89 ± 4% (mean ± SD, *n* = 3 mice). An anomaly dataset was constructed to simulate the drift of the recording using spikes gathered from a fourth independent mouse to test the drift detection ability of the autoencoder. The results showed that the mean squared error (MSE) between reconstructed and original waveforms (Fig. 5d) was higher for spikes detected from simulated drifting neurons, compared to the stable neurons used in the training and testing dataset. This significant difference allowed for drift detection based on reconstruction accuracy. A threshold could be used to distinguish the testing spikes and drifting spikes, which eliminated the majority of spikes from the drifting dataset (83 ± 12%, mean ± SD, *n* = 3 mice) and kept most spikes from the testing dataset (86 ± 7%, mean ± SD, *n* = 3 mice, Extended Data Fig. 8f). By visualizing the autoencoders’ embedding space in the bottleneck (Fig. 5e), cluster embeddings of the neuron spike waveforms from the same mouse showed higher separability than drifting spikes from a different mouse (Extended Data Fig. 8g-k). Furthermore, the training manifold convex hull was used as a stability verification tool for spike processing to quantify the within-boundary subset of testing dataset spikes (Fig. 5f and Extended Data Fig. 8h-k). Low amounts (8 ± 11%, mean ± SD, *n* = 3 mice) of out-of-manifold testing spike embeddings further demonstrated recording stability as a similarity in waveform shape between the first and last recording months, meaning the later spikes’ embeddings lay within the embedding space created by the first recorded spikes (Fig. 5f and Extended Data Fig. 8h-k). Collectively, high classification accuracy and low out-of-manifold percentage confirmed the long-term stable single-unit spike detected from the same neurons over time. This result also suggests that the stable recording data can be used to build the ML model to perform automated spike sorting based on the first few months of recording from a given mouse by capturing single-neuron waveform salient characteristics used as the input to the classification head. Additionally, the ML model successfully detected the spikes from a different mouse, highlighting the model’s ability to detect the drift of recording.

**Fig. 5.**
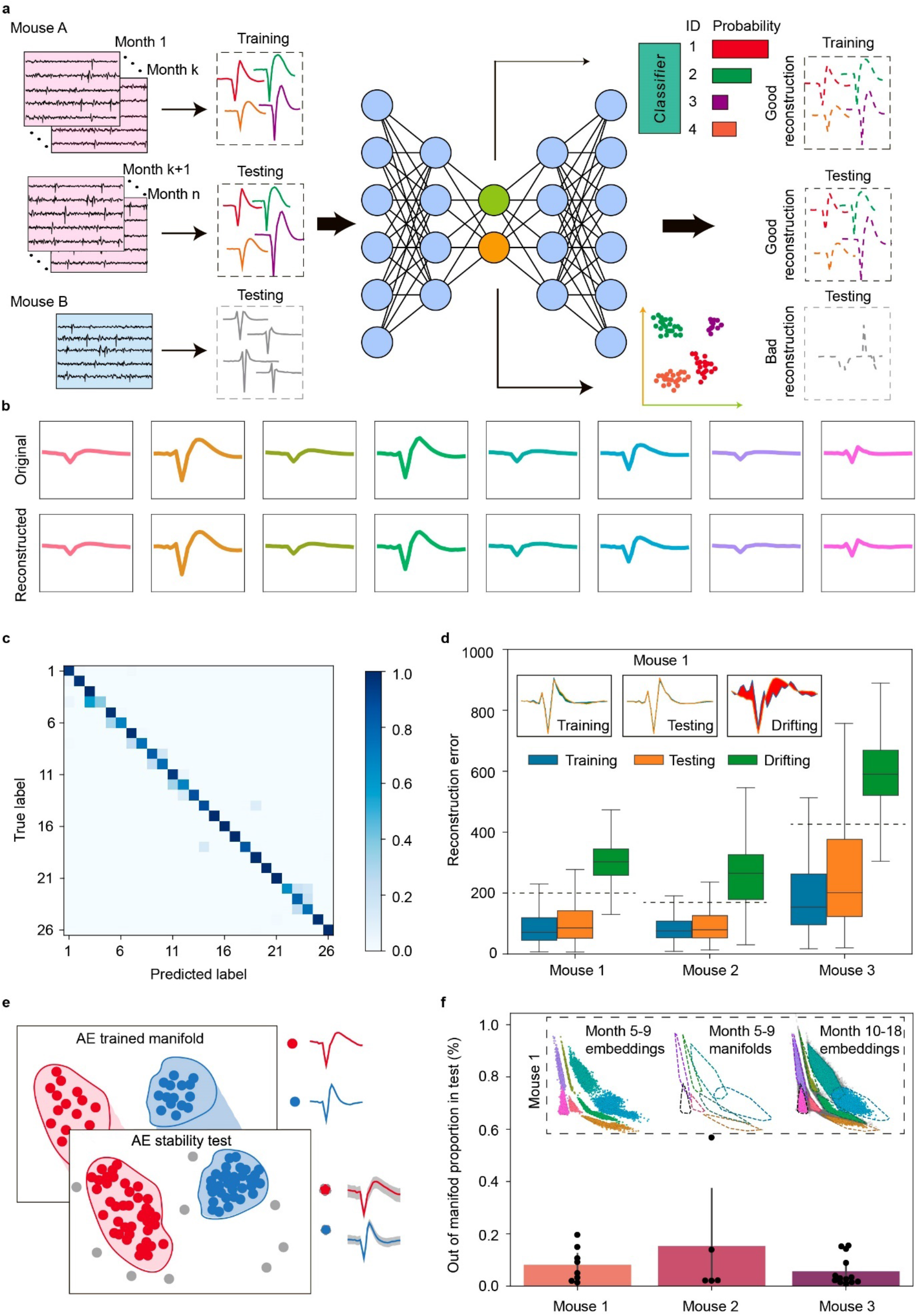
Long-term stable single-cell recording enabled autoencoder-based spike processing. **a**, Schematics of an autoencoder-based model which consists of an encoder, a classification head, and a decoder. Single-unit action potential waveforms are first extracted from the recording voltage traces and encoded with channel number to a lower-dimensional latent space. The latent representation is then used for classification and decoding. Spikes from the first 6 recording months including 5,490 spikes were used for training. The subsequent 8 months of data including 158,317 spikes were used for testing. Spikes from different groups of neurons collected from a different mouse were also used as simulated drifting data for testing. **b**, Original and reconstructed average representative single-unit waveforms from the testing data. Original waveforms are colored by neuron labels. Reconstructed waveforms are colored by the classification output. **c**, Confusion matrix comparing testing dataset ground-truth neuron labels with autoencoder classification output across all three mice. **d**, Boxplot of median and quartile range of mean-squared error (MSE) distribution for each of the training, testing, and drift datasets. Whiskers denote 1.5× interquartile range. Dashed lines indicate thresholds corresponding to mean + SD of mouse-specific MSE training distribution, eliminating 83 ± 4 % of training and 86 ± 7 % of testing (Extended Data Fig. 8f). Inset: representative average waveforms and their corresponding reconstructions by the mouse-specifically trained autoencoder. The colored areas correspond to the difference between the reconstructed and original waveform. **e**, Schematics showing the autoencoder can be used to quantify the stability of the recording. The trained manifold (highlighted by red and blue) was used to test the stability of the recording by quantifying the percentage of the neuron single-unit action potentials from the testing data falling inside the training manifold. **f**, Bar plot illustrating the percentage of the testing dataset for each neuron falling outside of the training manifold as per the process illustrated in (**e**). Data represented mean ± SD, individual data points are overlaid. Low levels of detected noise suggest recording stability through train-test waveform resemblance. Inset: applying noise rejection process to mouse 1 data, in order: training embeddings, training manifold boundaries, testing embeddings with out-of-manifold spikes colored in grey.

### Entire adult life study of brain aging at the single-neuron level

The adult mice life recording offers an opportunity to observe aging-associated electrical behavior changes at single-neuron resolution. We performed both qualitative and quantitative analyses of aging-associated changes at the single-neuron level over the mouse adult life. Analysis of clusters using PCA showed the stability of spikes from a group of neurons with largely overlapping clusters (purple and red, respectively) from each electrode, while the other spikes (green and blue) from the same electrode showed a slight change over the time course of recording (Fig. 6a). We quantitatively assessed the multivariate spread of cluster centroid positions by comparing average position shifts between consecutive cluster centroid positions (Fig. 6a, Methods) to average cluster distribution spreads. This assessment supported the stability (0.47 ơ_purple_, 0.58 ơ_red_) and variability (1.96 ơ_green_, 2.63 ơ_blue_) described previously. We analyzed the time-evolution of average neuron waveforms in a representative 3D feature space while calculating trajectories with a B-spline interpolation of successive positions (Fig. 6b, Methods). Non-correlated features were chosen after performing correlation analysis (duration, peak-trough ratio, and repolarization slope in Extended Data Fig. 9a). 21% of neurons (*i.e.,* green and blue) showed a clear trajectories trend while others (*i.e.,* purple and red) remained the same over the aging of the mice (Fig. 6b, Methods). We analyzed the time-evolution of the UMAP embeddings of spike waveforms (Fig. 6c, Extended Data Fig. 9b) in real time and ML-defined pseudotime^47^. To study the continuous and gradual transition of the neuron waveforms instead of the discrete real time label, we constructed a pseudo-temporal path termed as pseudotime to order spikes in the latent space using *monocle3*^47^, an ML tool originally for exploring the dynamics of gene expression within cell types and trajectories over time (Methods). The pseudotime of stable neurons (red and purple) remained the same value as the real time varied, which further validated the stability in the waveform during the aging-long recording. Similarly, the pseudotime of previously defined slow aging-associated neurons increased as the mouse got old. These qualitative and quantitative results suggest that the long-term stable recording from these open mesh nanoelectronics can track the aging-related electrical activity evolution from the same or same group of neurons in mice at single-cell level.

**Fig. 6.**
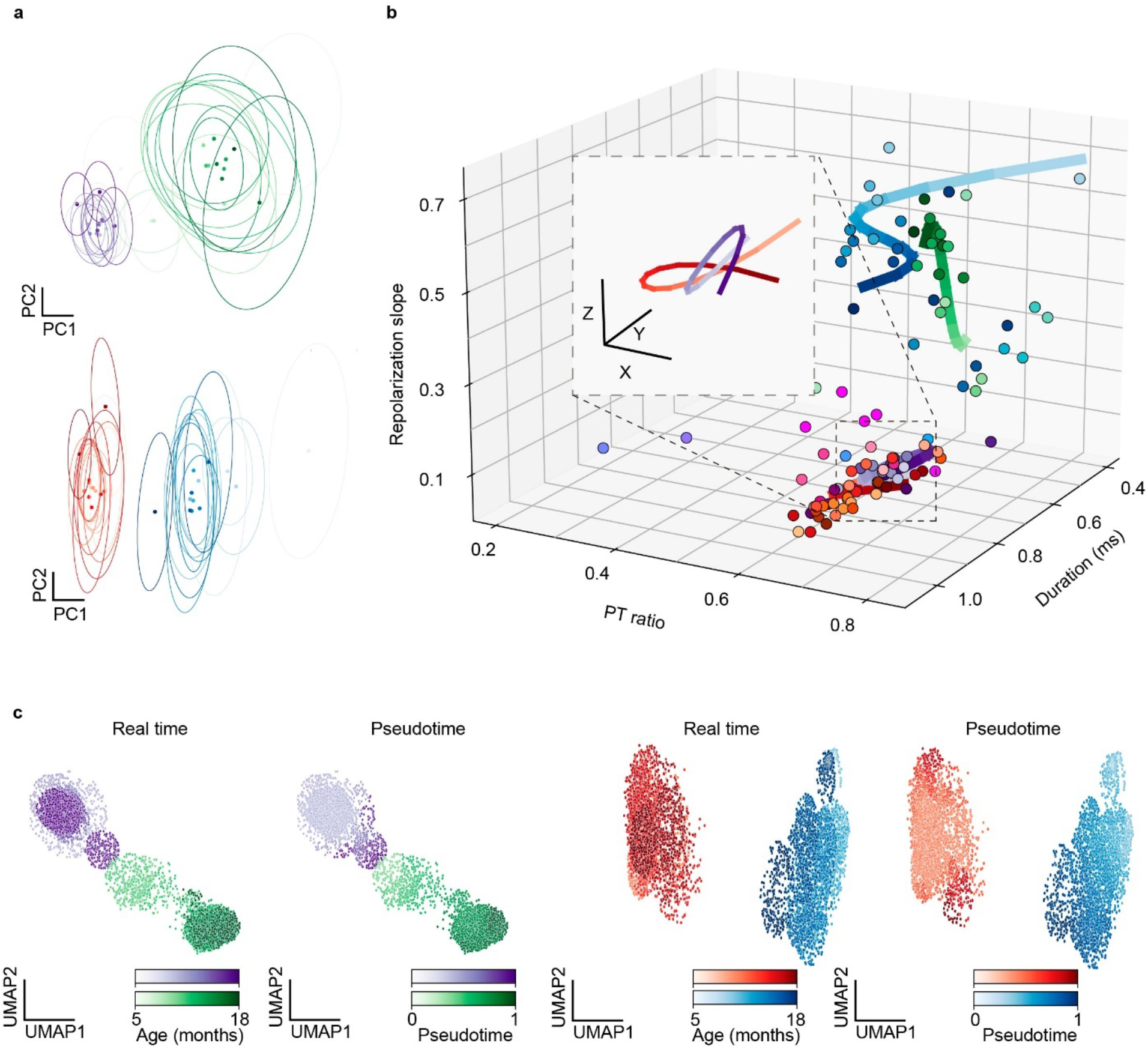
Adult life study of brain aging at the single-neuron level. **a**, Principal component analysis (PCA) of representative waveforms of neurons showing stable (purple and red) and progressive changing (green and blue) electrophysiological behaviors. Dots and ovals represent the centers and ±2 ơ of PC clusters, respectively. Average cluster centroid successive position shifts: 0.47 ơ_purple_, 0.58 ơ_red_, 1.96 ơ_green_, 2.63 ơ_blue_. Low values indicate stable cluster centroid position across time relative to single cluster whereas high values suggest time-evolution of corresponding cluster centroids and associated single-unit action potential waveforms. In both cases, a single electrode channel tracked stable (< 1ơ) and time-evolving single-unit action potential waveform. **b**, Trajectory analysis conducted with a B-spline interpolation of representative waveform features over the time course of implantation for the same neurons. Dots with the same color represented mean values of features calculated by the waveforms associated with the same color-coded neurons from data shown in (**a**). Trajectories highlight the representative neurons shown in (**a**) showing slow progressive and aging-associated electrophysiological properties. Inset: zoom-in view of purple and red trajectory. Scale bar, x = 0.01, y = 0.025, z = 0.01. **c**, Comparison of the electrophysiology Monocle3-based pseudotime and real time evolution over the adult life of mice of overall UMAP representation of representative individual single-unit action potential waveforms. Pairs of neurons recorded by the same electrode are compared.

## Conclusion

We demonstrated that the ultra-thin shuttle monolithically integrated mesh nanoelectronics can be implanted across multiple brain regions with an open mesh structure with minimal tissue damage. The open mesh structure is interwoven with the neural network in the brain of the animal, enabling immune response-free implantation and long-term stable 3D electrode-to-neuron integration. This structural stability allowed us to track the activity of the same neuron over the entire adult life of mice until natural death as supported by our extensive statistical data analyses showing stable impedance, waveform, firing dynamics, and recording performance, something not achieved by other *state-of-the-art* electrodes. We leveraged the high recording stability of this method, capable of successfully training an autoencoder using the first month’s recording, which further validate the stability of recording. Combining the stable recording and autoencoder, we can automate spike processing, sorting and stability verification on the remaining months’ recording. The unique ability to successfully track individual neurons in a chronically stable manner over such a long timespan provides a continuous view of aging-associated changes in neural activity. Combining the evolution of spiking activity at both real time and ML-calculated pseudotime, we observed potential aging-associated waveform changes at the single-neuron level. We believe long-term stable tracking of single neuron activity patterns across a stably recorded population of cells combined with automated data analysis tools will open new opportunities for the next-generation brain-machine interface and bioelectronic medicine. This technology also promises to inform our understanding of many long-term processes, including development, learning, recovery from injury, neurodegeneration and age-related cognitive decline. In the future, we envision that further integration of stretchability into our current device design, which can further adapt to the large volume change during early brain development, could further allow us to achieve the long-term stable recording over the entire lifespan of animals.

## Methods

### 1. Fabrication of ultra-thin shuttle monolithically integrated mesh nanoelectronics

(1) Cleaning a silicon wafer grown with thermal oxide (500-nm thickness) with acetone, isopropyl alcohol, and deionized water. (2) Depositing 100-nm-thick nickel (Ni) using a Sharon thermal evaporator as a sacrificial layer. (3) Spin-coating SU-8 precursor (SU-8 2000.5, MicroChem) at 3000 rpm, which was pre-baked at (65 °C, 95 °C) for 2 min each, exposed to 365 nm ultra-violet (UV) for 200 mJ/cm^2^, post-baked at (65 °C, 95 °C) for 2 min each, developed using SU-8 developer (MicroChem) for 60 s, and hard-baked at 180 °C for 40 min to define mesh SU-8 patterns for bottom encapsulation. (4) Spin-coating LOR3A photoresist (MicroChem) at 4000 rpm, followed by pre-baking at 180 °C for 5 min; spin-coating S1805 photoresist (MicroChem) at 4000 rpm, followed by pre-backing at 115 °C for 1 min; the sample was then exposed to 405 nm UV for 40 mJ/cm^2^, and developed using CD-26 developer (Microposit) for 70 s to define interconnects patterns. (5) Depositing 5/70/5-nm-thick chromium/gold/chromium (Cr/Au/Cr) by a Denton electron-beam evaporator, followed by a standard lift-off procedure in remover PG (MicroChem) to define the Au interconnects. (6) Repeating Step (4) to define electrode array patterns in LOR3A/S1805 bilayer photoresists. (7) Depositing 5/50-nm-thick chromium/platinum (Cr/Pt) by a Denton electron-beam evaporator, followed by a standard lift-off procedure in remover PG (MicroChem) to define the electrode array. (8) Repeating Step (3) for top SU-8 encapsulation. (9) Spin-coating SU-8 precursor (SU-8 2025, MicroChem) at 4000 rpm, which was pre-baked at 65 °C for 2 min and 95 °C for 8 min, exposed to 365 nm ultra-violet (UV) for 200 mJ/cm^2^, post-baked at 65 °C for 2 min and 95 °C for 6 min, developed using SU-8 developer (MicroChem) for 6 min, and hard-baked at 180 °C for 1 hour to define SU-8 anchors patterns to connect the mesh and SU-8 shuttle. (10) Spin-coating 20 wt% dextran solution at 1000 rpm for 20 s. which was at 80 °C for 1 min and 180 °C for 30 min. (11) Spin-coating SU-8 precursor (SU-8 2025, MicroChem) at 3000 rpm, which was pre-baked at 65 °C for 2 min and 95 °C for 8 min, exposed to 365 nm ultra-violet (UV) for 200 mJ/cm^2^, post-baked at 65 °C for 2 min and 95 °C for 6 min, developed using SU-8 developer (MicroChem) for 6 min, and hard-baked at 180 °C for 1 hour to define the SU-8 shuttle pattern. (12) Cleaning the input/output with water and soldering a 32-channel flexible flat cable (Molex) onto the input/output pads using a flip-chip bonder (Finetech Fineplacer). (13) Soaking the mesh nanoelectronics in nickel etchant for 2 to 4 hours to completely release the mesh nanoelectronics from the substrate wafer. (14) Rinsing the mesh nanoelectronics with deionized water and PBS three times each. (15) Dicing the substrate to the desired length. Dip-coating 10 wt% PEG solution to attach the mesh nanoelectronics and SU-8 polymer shuttle. The monolithically integrated mesh nanoelectronics was allowed to dry in the air. (16) After cutting the anchor, the monolithically integrated mesh nanoelectronics was ready for implantation.

### 2. Bending stiffness calculations

We estimated and compared the bending stiffness values (D) of mesh and thin-film nanoelectronics using a beam model. The bending stiffness of mesh nanoelectronics with three-layer polymer/mesh/polymer structure can be calculated as

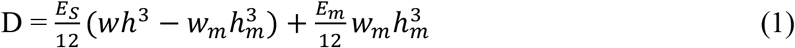

where E_s_ and E_m_ are the young’s moduli of SU8 and gold, 2 and 79 GPa, respectively, *w* is the width of SU-8 scaffolds, *w_m_* is the width of gold interconnects, *h_mesh_* = 0.91 μm and *h_film_* = 0.92 μm is the measured total thickness of SU-8, *h_metal_* = 80 nm is the thickness of Cr/Au interconnects. The calculated bending stiffness of mesh and thin-film nanoelectronics,1.26×10^−15^ N·m^2^ and 39.8 × 10^−15^ N·m^2^, respectively.

### 3. Brain implantation

All the animal experiments were approved by the Institutional Animal Care and Use Committee of Harvard University. The implantations were carried out on the male C57BJ/6 mice (∼16 weeks of age). The animals were housed in a regular 12 h/12 h light/dark cycle. Animals were anesthetized with 2-3% isoflurane and maintained under anesthesia with 0.75–1% isoflurane during the intracranial implantation surgery. Stainless-steel screws were implanted in the cerebellum and used as ground electrodes. A craniotomy (2 × 2 mm^2^) was performed on the brain, and the cortical surface was exposed upon removal of the dura mater. The mesh nanoelectronics with the releasable shuttle was attached onto a micromanipulator on the stereotaxic frame. The micromanipulator was manually controlled to insert the nanoelectronics into the mouse brain at the targeted depth at the tip. Sterile PBS was applied on the rear end of the nanoelectronics to dissolve the PEG/dextran and release the SU-8 shuttle from mesh nanoelectronics. After PEG/dextran was fully dissolved on both ends, the SU-8 shuttle was extracted with the second manipulator, leaving ultra-flexible mesh nanoelectronics implanted at the target positions. Craniotomies were sealed with a silicone elastomer (World Precision Instruments, USA). Ceramic bone anchor screws, together with dental methacrylate, were used to fix the FFC and electrode set onto the mice’s skull.

### 4. Immunohistochemistry

The following procedures were performed according to our previous reports^18,48^

#### Histology sample preparation

At each time point (2-, 6-, 12-week and 1-year post-implantation), mice were anesthetized with 40-50 mg/kg sodium pentobarbital and then transcardially perfused with ∼40 ml PBS pre-wash, and ∼40 ml 4 % paraformaldehyde (PFA) in PBS, followed by decapitation. The scalp skin was removed, and the exposed skull/dental cement were ground for 10–20 min at 15,000 r.p.m. using a high-speed micro motor tool. The brain with the mesh/thin film nanoelectronics undisturbed was removed from the cranium and postfixed in PFA for 24 h at 4 °C. The brain was transferred to incrementally increasing sucrose solutions (10–30%, w/v) until sunk to the bottom for the thin tissue preparation.

#### Immunohistochemical staining of 20-μm thin tissue

After cryostat sectioning, brain slices were incubated PBST (1 × PBS with 0.2% Triton X-100, Thermo Fisher Scientific) for 30 min, and then blocked with 5 % (w/v) normal donkey serum for 2 hours. After three rinses with PBST for 30 min each, slices were then incubated at 4 °C overnight in the primary antibodies: chicken anti-glial fibrillary protein GFAP (targeting astrocytes, 1:200, Abcam #ab4674, USA), goat anti-ionized calcium binding adaptor molecule 1 (Iba1) (targeting microglia, 1:100, Abcam #ab5076, USA), and rabbit anti-neuronal nuclear NeuN (targeting nuclei of neurons, 1:200, Abcam #ab177487, USA), followed by the slices being washed three times for 30 min each with PBST. Slices were then incubated in a secondary antibody solution at room temperature for 2 h with protection from light (1:500, Alexa Fluor 647 donkey anti-chicken, Jackson Immunoresearch, USA; 1:500, Alexa Fluor 594 donkey anti-goat; 1:500, Alexa Fluor 488 donkey anti-rabbit, Invitrogen, USA). After being washed three times for 30 min each with PBST, brain slices were also stained by incubating with 4’,6-diamidino-2-phenylindole (DAPI, Sigma-Aldrich, USA) to mark all cell nuclei for 30 min. After being washed, slices were mounted on glass slides with coverslips using Prolong Gold (Invitrogen, USA) mounting media. The slides remained covered (protected from light) at room temperature, allowing for 12 h of clearance before imaging.

#### Tissue clearing and staining for thick tissues

After vibratome sectioning, brain slices were placed in 1 × PBS containing 4% (w/v) acrylamide (Sigma-Aldrich) and 0.25% (w/v) VA-044 thermal polymerization initiator (Fisher Scientific) at 4 °C for 3 days. The solution was replaced with fresh solution immediately before placing the brain slices in X-CLARITY polymerization system (Logos Biosystems) for 3 h at 37 °C. After polymerization, any remaining gel from the tissue surface was removed and the slices were rinsed with PBST before placing them in electrophoretic tissue clearing solution (Logos Biosystems) at 37 °C for 3–5 days until the samples were translucent. Brain slices were incubated with PBST overnight, followed by three washes with PBST, and then blocked with 5 % (w/v) normal donkey serum for 2 days. After three rinses with PBST, slices were then incubated at 4 °C for 5-7 days in the primary antibodies containing: chicken anti-glial fibrillary acidic protein (GFAP) (targeting astrocytes, 1:200, Abcam #ab4674, USA) and/or and NeuN (targeting nuclei of neurons, 1:200, Abcam #ab177487, USA), followed by the slices being washed three times with PBST. Slices were then incubated in a secondary antibody solution at 4 °C for 5-7 days with protection from light (1:500, Alexa Fluor 647 donkey anti-chicken, Jackson Immunoresearch, USA; 1:500 and/or Alexa Fluor 488 donkey anti-rabbit, Invitrogen, USA). After being washed three times with PBST, brain slices were also stained by incubating with 4’,6-diamidino-2-phenylindole (DAPI, Sigma-Aldrich, USA) to mark all cell nuclei for 2 days. Brain slices were glued at their edge to the bottom of Petri dishes with 1% (w/v) agarose in optical clearing solution (Lifecanvas Technologies) 24 h before microscopy imaging.

### 5. Microscope imaging and image data analysis

Confocal fluorescent images were acquired using a Leica SP8 confocal system (Leica, USA). Images were collected using a 25×, 0.95 NA water-immersion or 40×, 1.3 NA oil-immersion objective lens. 488 nm, 591 nm, and 633 nm lasers as the excitation sources for Alexa Fluor 488, Alexa Fluor 594, and Alexa Fluor 647, respectively. Standard TIFF files were exported and colorized using LAS X Software. ImageJ software and custom code were used for image analysis. The distance of each pixel to mesh/film nanoelectronics was defined as its shortest distance from the mesh/film boundary. Baseline fluorescence intensity is defined as the average fluorescence intensity of all pixels 525–550 μm away from the boundary. Intensity values with distances binned over an interval of 25 μm were averaged and normalized against the baseline intensity.

### 6. Electrophysical measurement

Mice with implanted mesh nanoelectronics and FFC connector were recorded chronically monthly using CerePlex Direct recording system (Blackrock Microsystem, USA), starting from 1-month post-implantation. Mice were anesthetized with 1 % isoflurane in medical-grade O_2_ or head-fixed during the measurement. Homemade PCB for connecting the FFC and head stage. Two animals are excluded from the long-term study (Fig. 4) since the poor body condition and connectors failed at an early stage. The electrophysiological recording was made with a 30-kHz sampling rate and a 60-Hz notch filter. Whiskers of interest were trimmed at ∼15 mm from the face and inserted into a polyimide tube fixed to the galvanometer system (PT-30K, SpaceLas, China) positioned ∼10 mm from the vibrissal pad to yield high-fidelity sensory stimuli. Stimulation was always delivered along the rostro-caudal axis. Voltage command and output for the actuator were programmed by Axon Digidata 1550B (Molecular Devices, USA).

### 7. Data analysis

#### Spike sorting and clustering

All recording data was analyzed offline. 6 min continuous recordings were used for analysis of each month. Spike detection was performed using the *WaveClus3* software package (https://github.com/csn-le/wave_clus). In brief, raw recording data was filtered using four poles Butterworth filters in the 300-3000 Hz frequency range before spikes were detected using an amplitude threshold 5 times the estimated standard deviation of the noise. After spike alignment, 30 data points with the sampling rate of 10 kHz were kept for each detected spike representing 3 ms. No normalization procedure was applied to the spike waveforms due to the stable nature of recorded amplitudes (Fig. 4f) and experiments (Extended Data Fig. 10a-f) showing normalization decreased putative neuron cluster separability as well as meaningfulness of PCA embeddings. Spike sorting results of *WaveClus3* obtained through superparamagnetic clustering were kept and used for comparison with the results of our chosen spike sorting approach. Additional quality metrics calculated for individual neurons demonstrate that the individual neurons were clearly defined (Extended Data Fig. 10g-i). Results reported in Figure 4 and 6 used clusters determined by *Leiden*^49^(https://github.com/vtraag/leidenalg) clustering performed on the graph constructed by Uniform Manifold Approximation and Projection (UMAP, https://github.com/lmcinnes/umap) for each individual month and over individual channels. Geometric considerations ensured that no neuron was being recorded by two separate channels. Cluster labels were then aligned by choosing a label matching scheme which minimized the mean-squared error between the average cluster waveforms of a given month and a chosen template.

#### Stability analysis of single-unit action potentials

After spike detection, sorting and clustering, stability analysis was performed. A custom computational pipeline for assessing the stability analysis, which was built using Python v3.9.4. Performing prior spike extraction using third-party software or using the pipeline to perform spike extraction and sorting directly in python are both possible.

UMAP analysis was used to confirm the recording stability. UMAP embeddings’ coordinates were calculated for all spikes of a given channel before plotting the corresponding points per recording month producing a visualization over time. The representations originating from different channels were plotted next to each other after manual curation for quality control to ensure only the best quality spikes were included in the analysis. Additionally (Fig. 6a), average cluster centroid position shifts were compared to average cluster distributions spread. Concretely, let us name *X* ∈ ℜ *^dx^*^2^ a vector of cluster centroid positions over d days in a given 2D (x, y) embedding. For each identified neuron, we calculated:

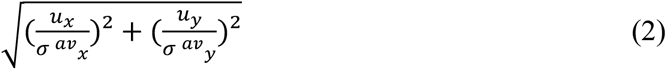

with 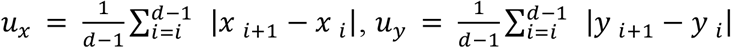 the average successive cluster centroid absolute position shifts along each axis, and 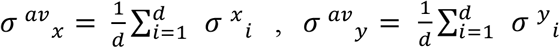 the average cluster distribution SD along each axis. This allowed us to compare centroid position shifts with average cluster SD along each axis to examine the embedded waveform stability. Results of correlation analysis using Pearson correlation coefficients and associated two-sided *t* tests were carried out using *scipy* v1.6.3 (http://www.scipy.org) stats module.

Cluster quality metrics shown in Figure 4d were obtained by calculating silhouette score using scikit-learn v0.24.2’s (http://scikit-learn.org) implementation over individual channels and months. L-ratio for an individual cluster was calculated as

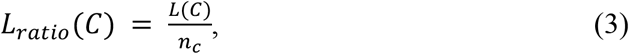

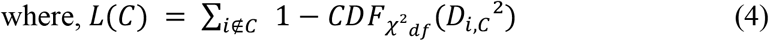

with *n_c_* the number of spikes belonging to cluster C, *CDF_x_^2^_df_* the cumulative distribution function of the *χ*^2^distribution, *df* = 2in our case and *D_i,c_*^2^ being the mahalanobis distance of spike i from the center of cluster C. Linear regressions were performed using scikit-learn. Within unit correlation analysis in Figure. 4c was calculated based on the Pearson product-moment correlation coefficient. The *corrcoef* function in *numpy* v1.18.5 (http://www.numpy.org) was used between all pairs of average cluster waveforms and recording days to provide the auto and cross correlation coefficients. Feature extraction was performed by using functions adapted from *AllenSDK* (https://github.com/AllenInstitute/AllenSDK).

#### Analysis of interspike intervals

The spiking times of each sorted neuron were used to calculate the interspike interval (ISI) histograms for individual months per cluster with a bin size of 2 ms.

#### Data analysis of whisker-stimulus provoked recording

For whisker responsive recording, firing timing of each detected single-unit spike was presented and aligned by trial number in the raster plot. For the peristimulus time histogram (PSTH), 1 ms bin size was used and the spike counts were accumulated from all trials within each recording session.

#### Autoencoder-based automatic spike processing

All autoencoders were built using *tensorflow 2.5* (https://www.tensorflow.org). The encoder consisted of two fully connected (FC) layers with 100 nodes each before the bottleneck 2D layer. Inputs to the network were concatenations of individual spike waveforms and their corresponding one-hot encoded channel (geometric considerations of the sparsity of the mesh recording device ensure that no single neuron can be recorded on two separate channels). From the bottleneck, the classification head, which consists of a single FC layer with *softmax* activation, outputs probabilities of belonging to a particular neuron class. The decoder’s symmetric structure with respect to the encoder allows for reconstruction of the original waveform. Activations for encoder and decoder layer were set as *LeakyReLU* with alpha = 0.3 as well as an added *L*_2_ regularizer on the encoder’s last layer, enforcing lower latent embedding values. Adam optimizer with default parameters was used. For a given dataset, *X* ∈ *R^nx^*^30^, *n* ∈ ℵof n spikes, each composed of 30 data points recorded on c channels, we denote the input batch to the autoencoder 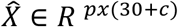. The autoencoder considered here is composed of an encoder, *f_e_*, decoder *f_d_* and classifying head *f_c_* parametrized by *θ* = (*θ_e_*, *θ_d_*, *θ_c_*). The network’s (slightly simplified, as the encoder’s last layer regularization is not explicit here) loss function over a batch of p spikes is thus:

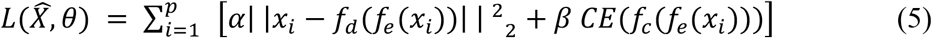

where CE is the usual categorical cross-entropy loss used for multi-class classification problems. The loss function is clearly defined by two components: reconstruction and classification, each with their own weight to regulate their overall contribution to the loss function (reconstruction error is also on a different scale than the cross-entropy loss so the relative weighting terms *α*and *β*are necessary for balancing contributions of the losses). Training the network consists of learning the correct parametrization *θ* (network weights) which minimizes the loss function over the training dataset. Early Stopping was applied on an evaluation dataset to avoid overfitting of the training data. One autoencoder was trained per mouse using a subset of that mouse’s first months recording data. The subset was chosen using conservative thresholds on the within-cluster centroid distance of UMAP embedded spike waveforms. This allows to train the autoencoder in a very short time on high-quality spikes without hurting classification results (Extended Data Fig. 8b). It also means the autoencoder’s training manifold can be used as a stability verification tool (Fig. 5f and Extended Data Fig. 8h-k). Indeed, waveforms recorded in the later parts of the recordings with similar shapes as the high-quality training waveforms will fall inside the training manifold.

#### Aging analysis at the single-neuron level

UMAP coordinates of the representation in Fig. 4 were used to visualize the different neuron trajectories shown in Figure 6 defined by their electrophysiological properties across the adult mouse life recording. Indeed, these UMAP embeddings were used to compute a trajectory graph with the corresponding “pseudo temporal” scale of evolution, with *Monocle 3* (https://cole-trapnell-lab.github.io/monocle3). The convex hull delimiting regions spanned by individual neuron time-evolution was calculated using *Scipy*; Principal Component Analysis (PCA) was used as another form of dimensionality reduction and was calculated per channel and month to view the evolution of spike waveforms per cluster (Extended Data Fig. 6b). Clusters obtained from the same channel were used to obtain cluster centroids per channel for individual recording months along with 2 SD confidence ellipses of corresponding covariance. Confidence ellipses were centered on cluster means while radii were calculated without explicit computation of eigenvalues by using Pearson correlation coefficients and equations (6) and (7):

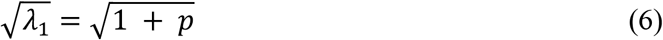

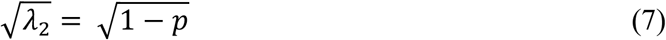

where *λ*_1_ and *λ*_2_ are the eigenvalues of the covariance matrix and p is the Pearson correlation coefficient. These equations hold for normalized data; thus, ellipses were scaled by twice the standard deviation along each axis to render the final plots.

Feature trajectory analysis was performed by fitting trajectories, from feature embeddings in a selected 3D feature space of mean feature values per cluster over real recording months. The features chosen for the representation were repolarization slope, peak trough ratio and duration. These were selected based on prior correlation analysis (Extended Data Fig. 9a) to make sure the trajectory was not a degenerate representation of feature evolution because of highly correlated features. Trajectory fittings were constructed using a B-spline interpolation on previously discussed points by first finding the parametric definition of the curve using *Scipy splrep* function before evaluating the spline using *splev* function. A cubic (*k* = 3) spline was used with s=2. as the smoothing condition in *splrep*.

## Acknowledgements

We acknowledge the discussion and assistance from all Liu Group members, Jane Salant, and Prof. Bence P. Ölveczky. We acknowledge the support from the Harvard University School of Engineering and Applied Sciences’ Startup fund and Harvard University Faculty of Arts and Sciences’ Dean’s Competitive Fund for Promising Scholarship.

## Author contributions

J.L. and S.Z. conceived and designed the experiments. S.Z., R. L, J. Lee. fabricated and characterized the electrodes. S.Z., Z.L. performed the brain implantation, *in vivo* recording and histology study. S.Z., X.T., S.P. conducted the data analysis. The manuscript was written by J.L., S.Z., X.T., and S.P.

## Extended Data Figures

**Extended Data Fig. 1.**
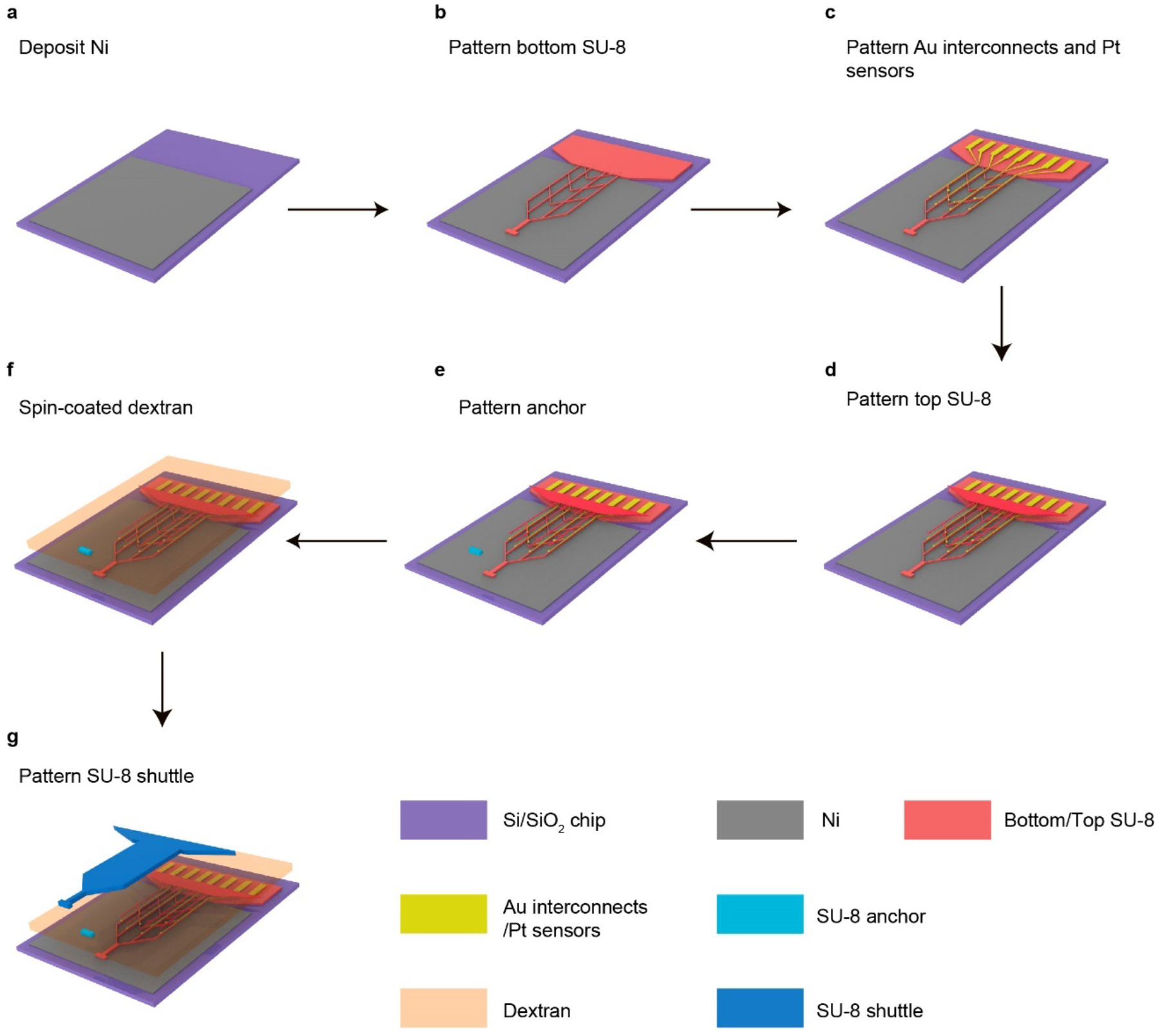
Schematics show the stepwise fabrication of monolithically integrated tissue-level flexible mesh nanoelectronics. **a**, A Ni sacrificial layer (grey) was defined by photolithography and deposited through thermal evaporation on the Si/SiO_2_ wafer (purple). **b**, SU-8 2000.5 bottom passivation layer (red) was defined by photolithography. **c**, Cr/Au interconnects and Pt microelectrodes (yellow) were sequentially defined by photolithography and deposited through electron beam (e-beam) evaporation on the top of the SU-8 passivation layer. **d**, SU-8 2000.5 top passivation was defined by photolithography (red). **e**, SU-8 2025 anchor was defined by photolithography (cyan). **f**, Dextran sacrificial layer (pink) was spin coated. **g**, SU-8 2025 shuttle was defined by photolithography (navy).

**Extended Data Fig. 2.**
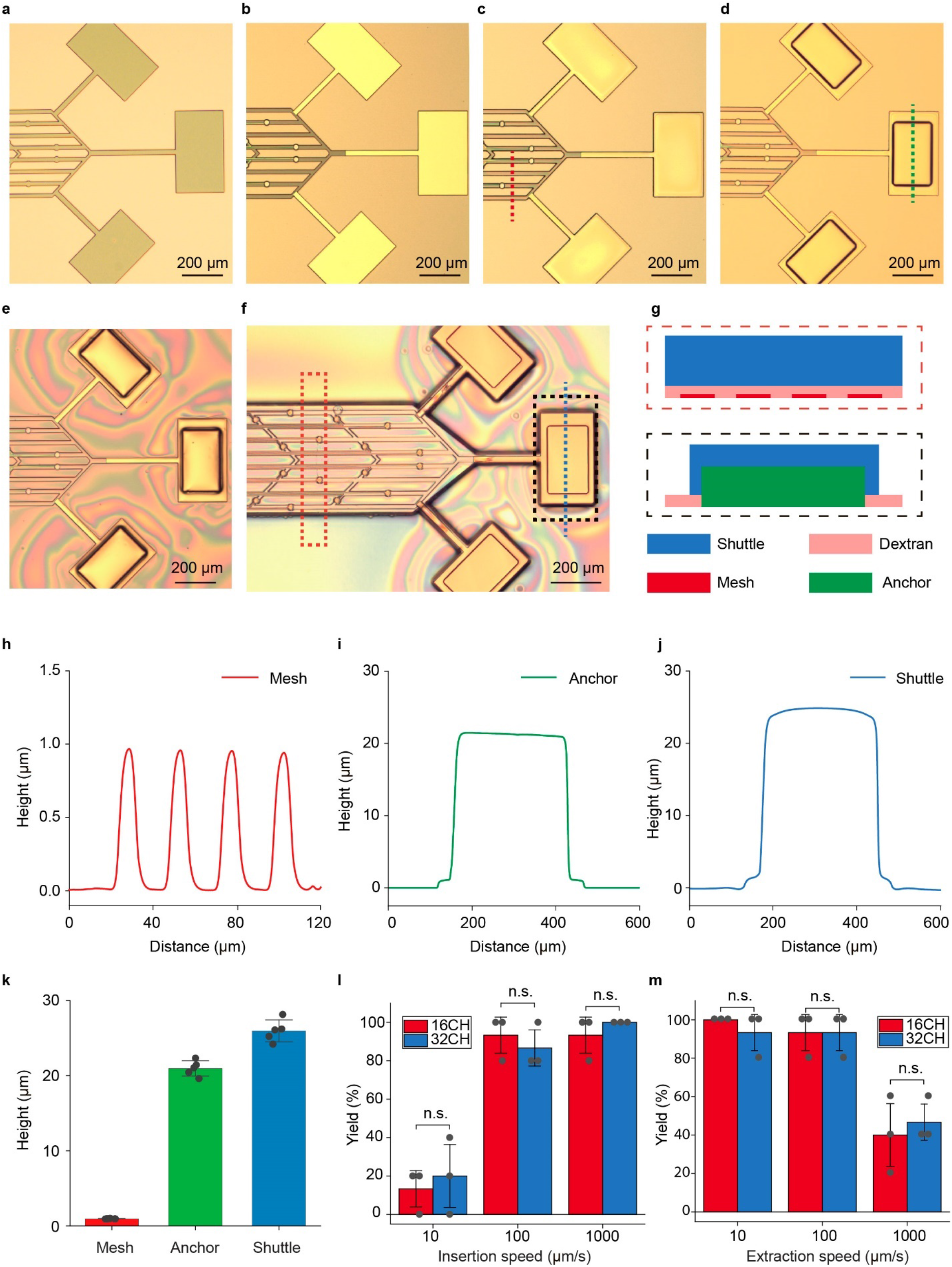
Anchor structures for controllable implantation. **a-f**, Optical images illustrating each step of the fabrication corresponding with Extended Data Fig. 1b-g, respectively. **g,** Schematics showing the cross-section of the monolithically integrated mesh nanoelectronics at the red and black dashed boxes highlighted regions in (**f**), respectively. **h-j,** Contact profilometer measurements along with the open mesh structure in (**c**, red dashed line), anchor structure in (**d**, green dashed line), and shuttle structure in (**f**, blue dashed line). **k,** Statistical summary of the thickness of the open mesh, anchor, and shuttle layer structures (*n* = 5). **l, m**, Insertion (**l**) and extraction (**m**) yield of 16-channel and 32-channel mesh nanoelectronics with different speeds (n.s.: not significant, two-tailed unpaired *t* test, *n* = 3 times experiment, each time include 5 insertion/extraction).

**Extended Data Fig. 3.**
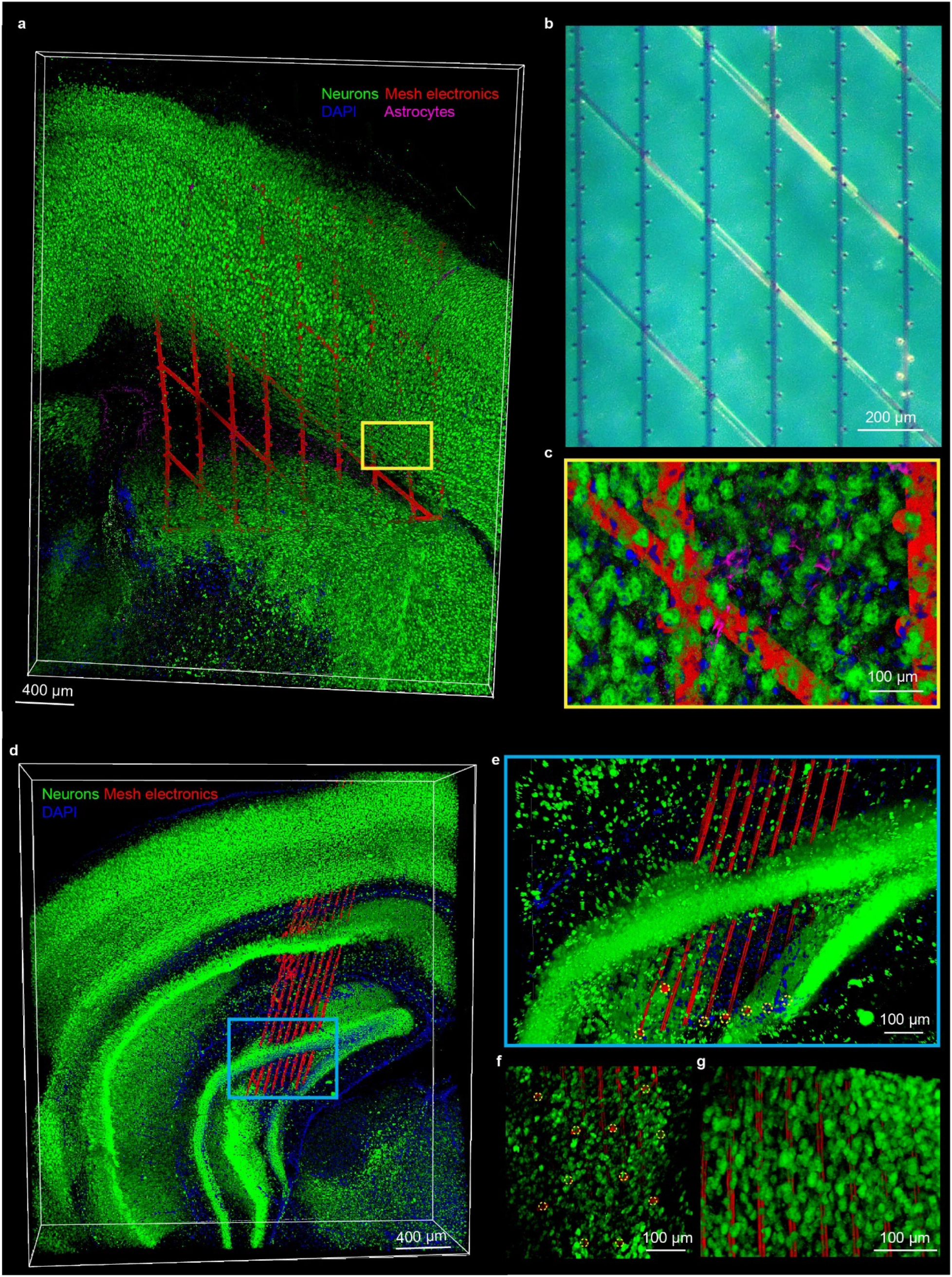
Independent replicates of mesh nanoelectronics with different sizes were implanted in mice brains. **a**, 3D reconstructed confocal fluorescence imaging of neuron (green), nuclei (blue), and astrocytes (purple) with 1024-channel mesh nanoelectronics (red) sustaining their open mesh structure across multiple brain regions at 6-week post-implantation. **b**, Representative photograph illustrates the high density, 1024-channel mesh nanoelectronics. **c**, Zoom-in views of the regions highlighted by yellow (**c**) box in (**a**). **d**, A 3D reconstructed interface of neurons (green), nuclei (blue) with shape maintained 16-channel mesh electronic (red) at 6-week post-implantation. The mesh electronic was across the cortex and hippocampus with a designed 30-degree angle corresponding to the dorsal-ventral direction. **e**, A zoom-in view of the hippocampus region highlighted by the cyan box in (**a**). **f**, **g**, Neuron interpenetration inside the subcellular electrode, individual electrodes are indicated by yellow dashed circles in (**e**) and (**f**) and opening mesh structure (**g**). These results show neuron interpenetration inside the opening mesh and minimal astrocyte increases at the surface and interior of mesh, and thus demonstrate the reproducibility of these results in Fig. 2.

**Extended Data Fig. 4.**
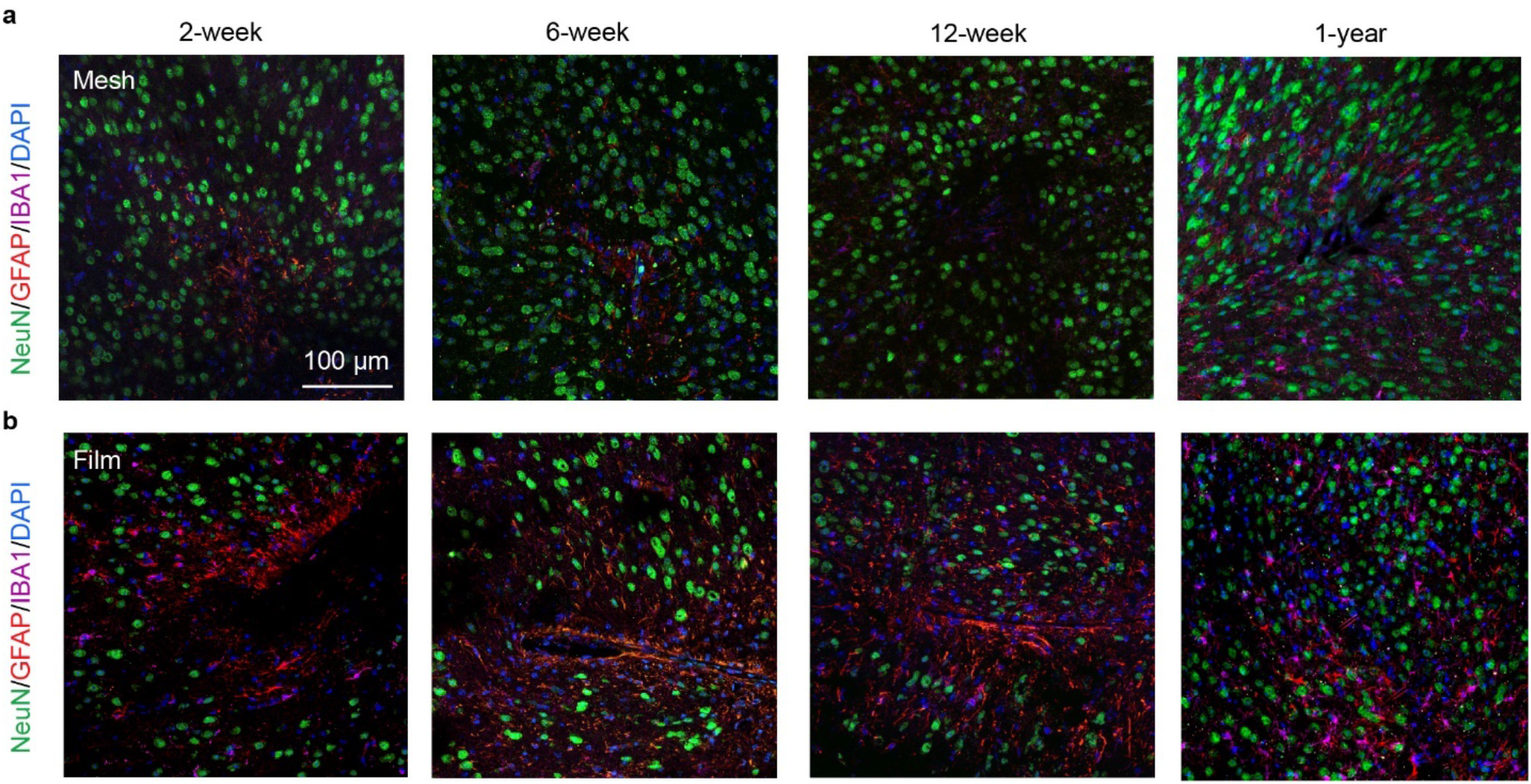
Time-dependent histology studies of brain tissue reaction to ∼1 μm-thick open mesh/thin-film nanoelectronics. **a**, **b**, Representative immunofluorescence images of brain tissue reaction following 2-week, 6-week, 12-week, and 1-year post-implantation of a mesh (**a**) and a thin-film (**b**) nanoelectronics from the contralateral hemisphere. The tissue was labeled for astrocytes (red), microglia (purple), neurons (green), and nuclei (blue). Time-dependent histology studies have been repeated on *n* = 5 independent samples for each time point, with statistical analyses shown in Fig. 2f-i.

**Extended Data Fig. 5.**
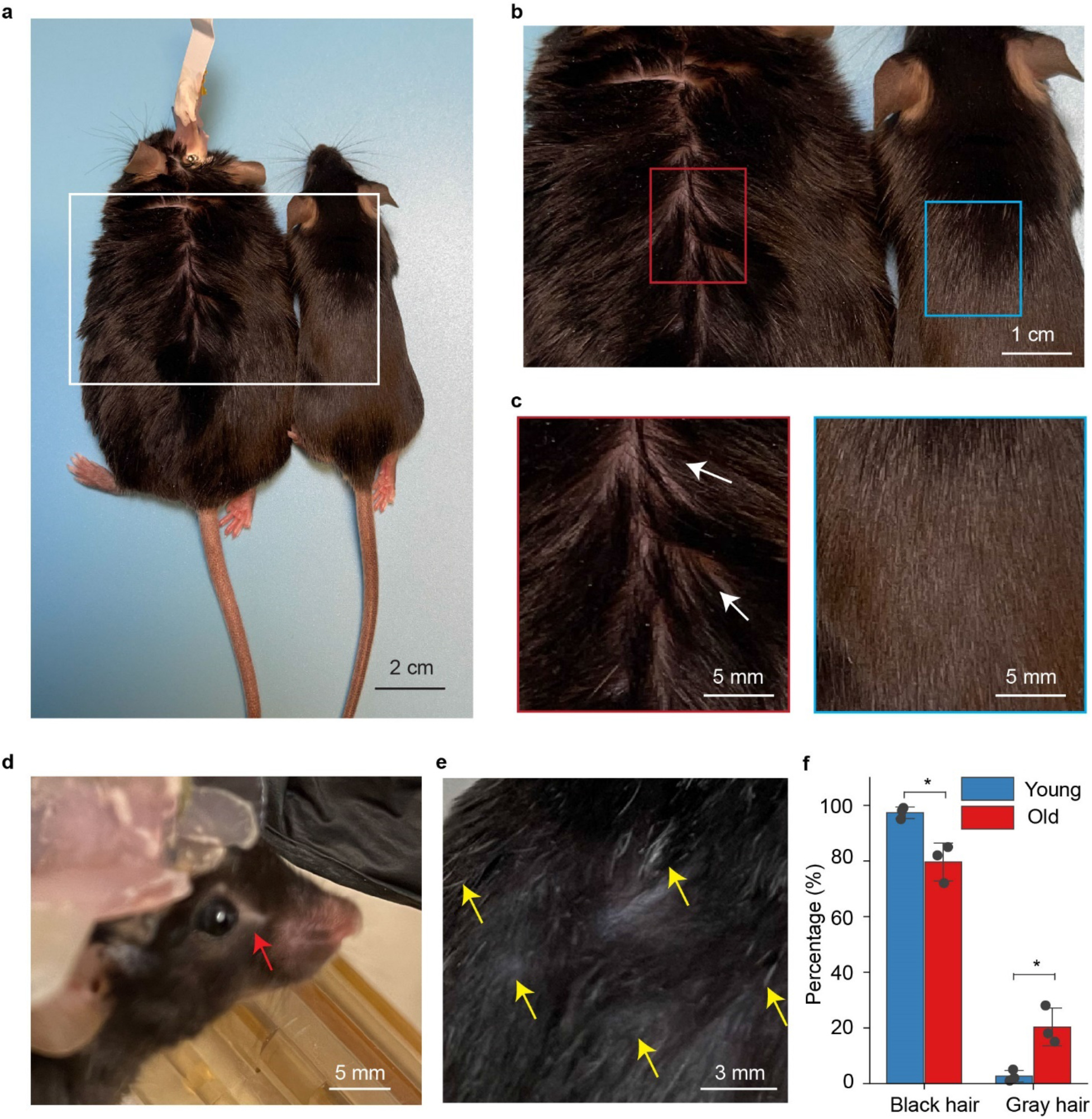
Aged mice characterization. **a**, Representative photograph showing the old mouse of weight gain (18 months) with tissue-like mesh nanoelectronics implant (left) compared with the mature adult mouse (5 months, right). **b**, Zoom-in views of the regions highlighted by white boxes in (**a**). **c**, Zoom-in views of the thinning hair (white arrows) of aged mouse and glossy brown fur of mature adult mouse highlighted by red and blue boxes in (**b**). **d-e**, Representative photograph showing the barbering around eyes (**d**, red arrow), grey and thinning fur in the dorsal back skin (**e**, yellow arrows) of the aged mouse (18 months) with mesh nanoelectronics implant. Statistical analysis reveals that significantly increased gray hairs in dorsal back skin in old-aged mice (*\*p* < 0.05, two-tailed unpaired *t* test, *n* = 3).

**Extended Data Fig. 6.**
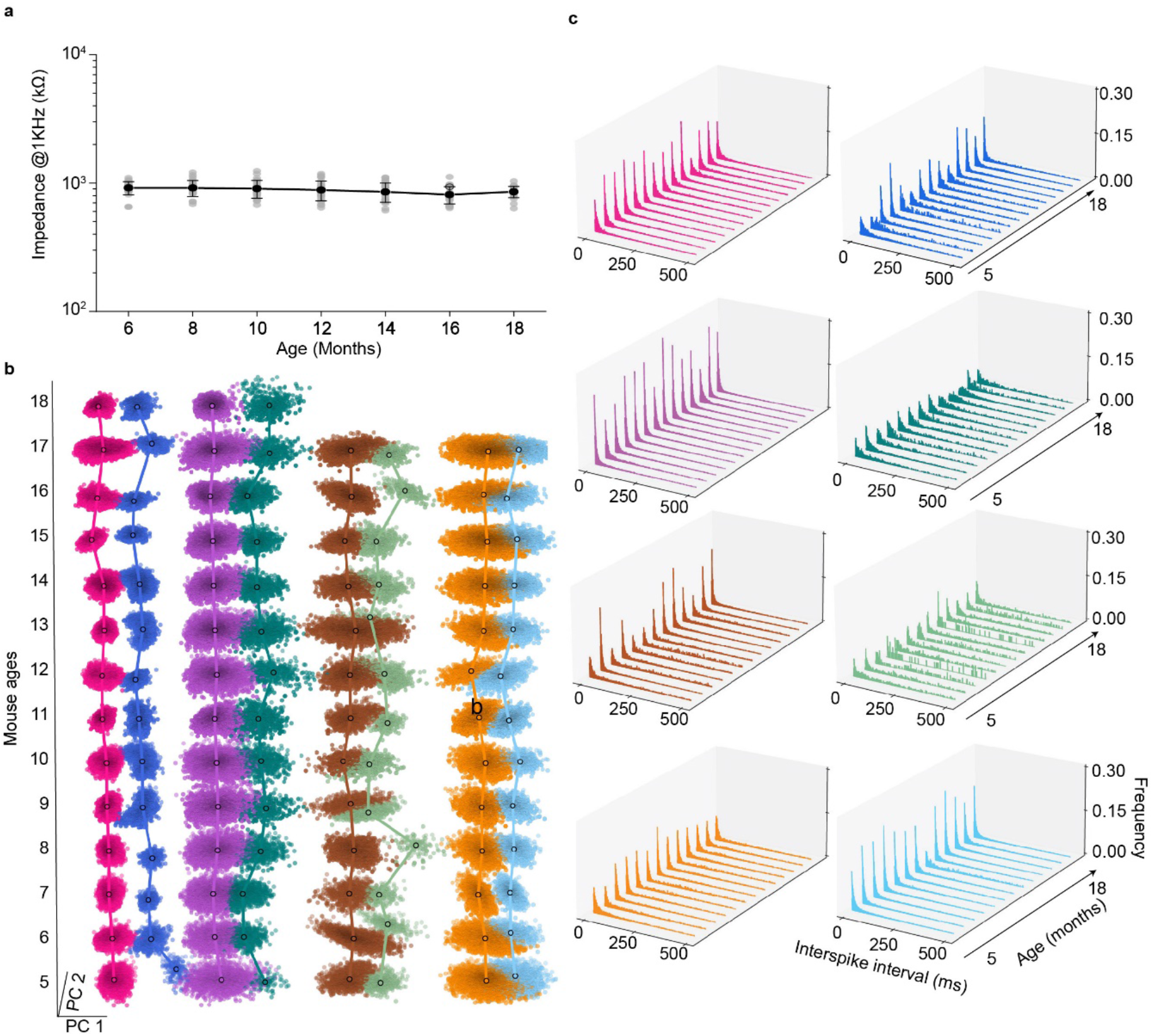
Long-term stable recording characterization. **a**, Time-dependent electrode interfacial impedance at 1 kHz measured by the Cereplex Direct (Blackrock Microsystems, USA). Data represented mean ± SD, individual data points are overlaid. The electrode interfacial impedances exhibited relatively constant values of 920.2 ± 107.2 kΩ vs. 857.2 ± 85.7 kΩ (mean ± SD, *n* = 30) at months 6 vs.18. **b**, Time evolution of representative single-unit spikes clustered by *Leiden* clustering. The x- and y-axes denote the first and second PC dimensions, respectively, and the z-axis denotes mouse age in months. Dimension-reduced clusters associated with a single unit are shown (Fig. 4a) using the same color. These data show stable clusters with nearly constant positions in the first and second principal component plane (PC1-PC2) over the entire recording period from 5 months to 18 months. **c**, Time evolution of interspike interval (ISI) histograms of representative neurons identified in Fig. 4 from 5 months to 18 months. The x- and y-axes denote the time between subsequent action potentials of spontaneous firing neuron, and mouse age in months, respectively, and the z-axis denotes frequency. Bin size, 2 ms.

**Extended Data Fig. 7.**
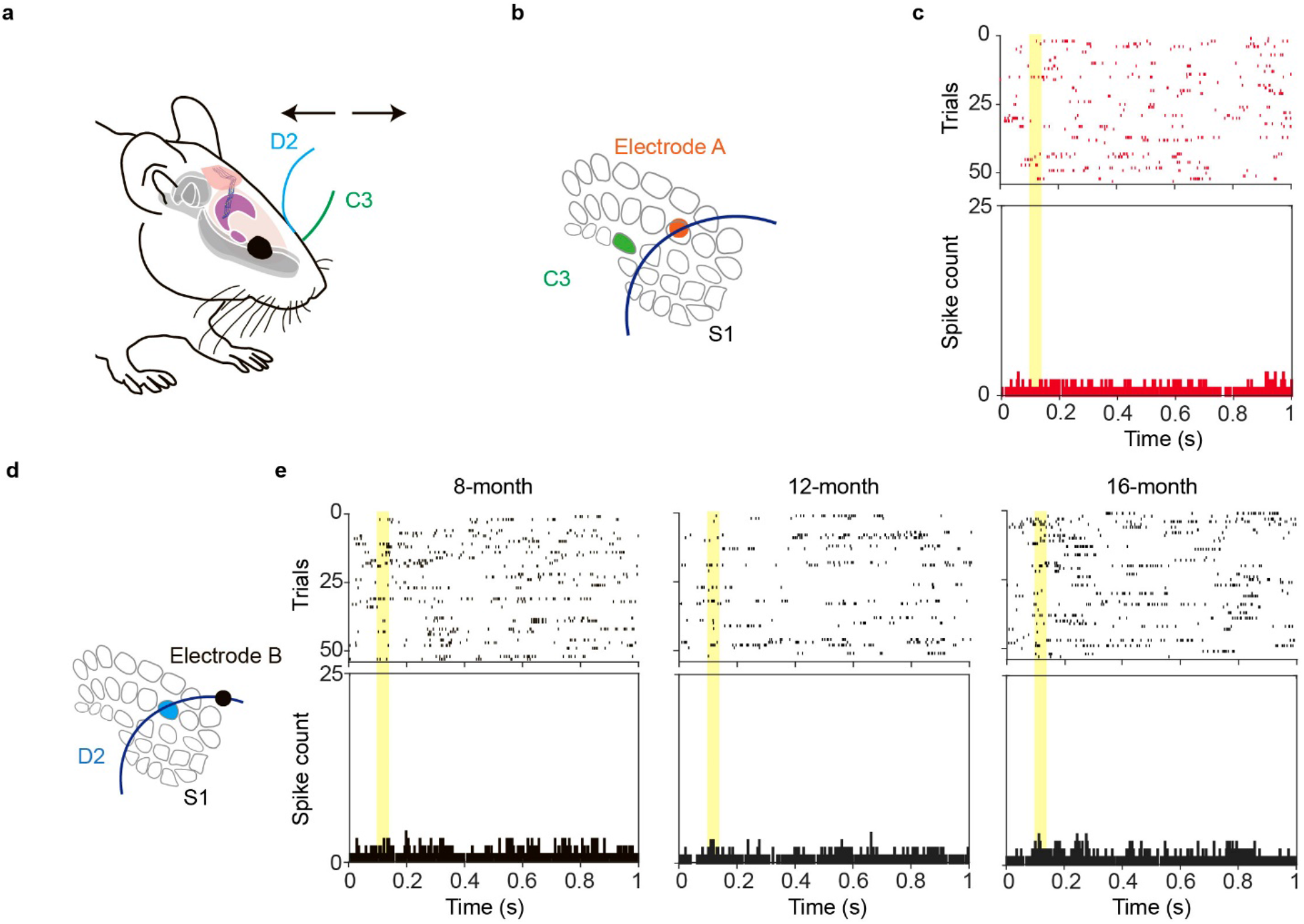
Validation of whisker-related activity using sham stimulation. **a**, Schematic diagram of whisker deflection. Individual vibrissa was deflected in the rostral-caudal plane using a computer-controlled piezoelectric bending actuator. D2 and C3 whiskers are labeled in blue and green, respectively. **b**, Schematic diagram of whisker barrel and electrode arrangement in S1. C3 whisker barrel and electrode A position is labeled in green and orange, respectively. **c**, Raster plot and peri-stimulus time histogram (PSTH, 1 ms bin size) of electrode A show no observable spiking activities when applied to the C3 whisker at 8 months. **d**, D2 whisker and electrode B position are labeled in blue and black in S1 barrel field, respectively. **e**, Raster plot and PSTH (1 ms bin size) of electrode B show no observable spiking activities when applied to the D2 whisker from 8 months to 16 months.

**Extended Data Fig. 8.**
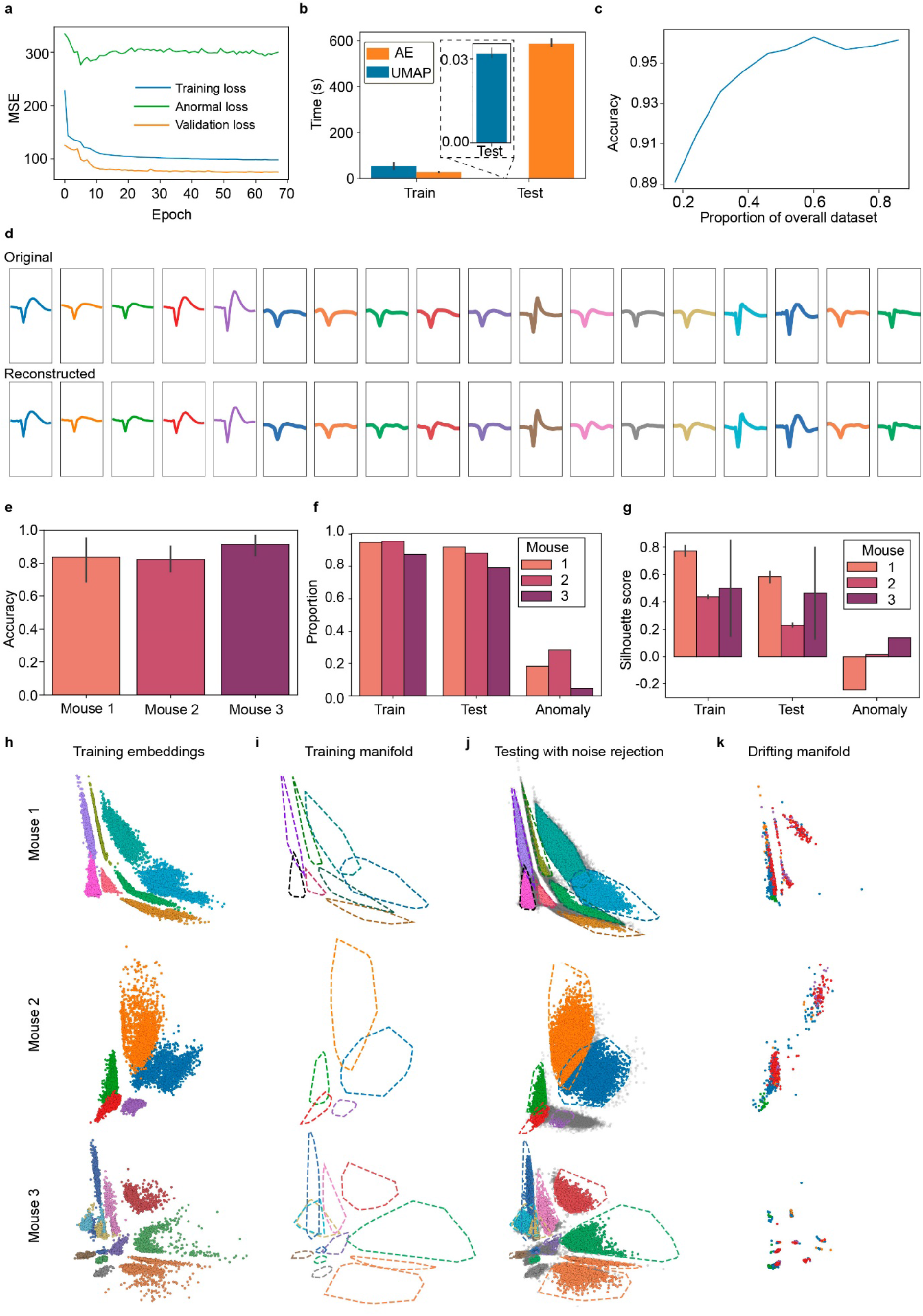
Autoencoder-based spike sorting and stability analysis. **a**, Representative training, validation, and drift loss curves obtained during the training process of one of the autoencoders. **b**, Time performance comparison of autoencoder vs computational pipeline allowing for dimensionality reduction and classification (UMAP and a Random Forest classifier trained on UMAP embeddings). Tests were performed using a single CPU with Intel i5, 8 cores @ 4.1GHz. The autoencoder has a significant time advantage for inference. **c,** Overall accuracy on the testing dataset as a function of the proportion of the overall dataset used for training the network. **d**, Original and reconstructed average neuron waveforms for the second and third mice data. Original waveforms on the top row are colored by within-mouse neuron labels (the first 5 neurons from the left are from mouse 2) while reconstructed waveforms on the bottom row are colored by autoencoder classifier predicted labels. Reconstruction and classification are near-perfect. **e**, Bar plot of per-mouse classification accuracy calculated for each neuron label class. **f**, Proportion of spikes kept for each dataset when using MSE-based thresholds for drift detection (Fig. 5d). **g**, Silhouette score calculated using autoencoder embeddings of training, testing and drift datasets with associated true neuron labels. The training and testing scores reflect the emergent latent space cluster separability for observed neurons while the anomalous scores highlight the poor separability of previously unseen embedded neuron waveforms. **h**, **i**, **j**, Stability verification process illustrated for all three mice: visualizing training latent space embeddings (dots colored by their true neuron label), calculating training manifold boundaries, and finally applying these boundaries to quantify the proportion of testing dataset spikes latent embeddings lying inside the predicted neuron’s training manifold. **k**, Visualization of the latent embeddings of drift spikes. Dots are colored by their true neuron labels; mixed colors within clusters showed poor ability to separate different neurons from the drift dataset, in accordance with (**g**).

**Extended Data Fig. 9.**
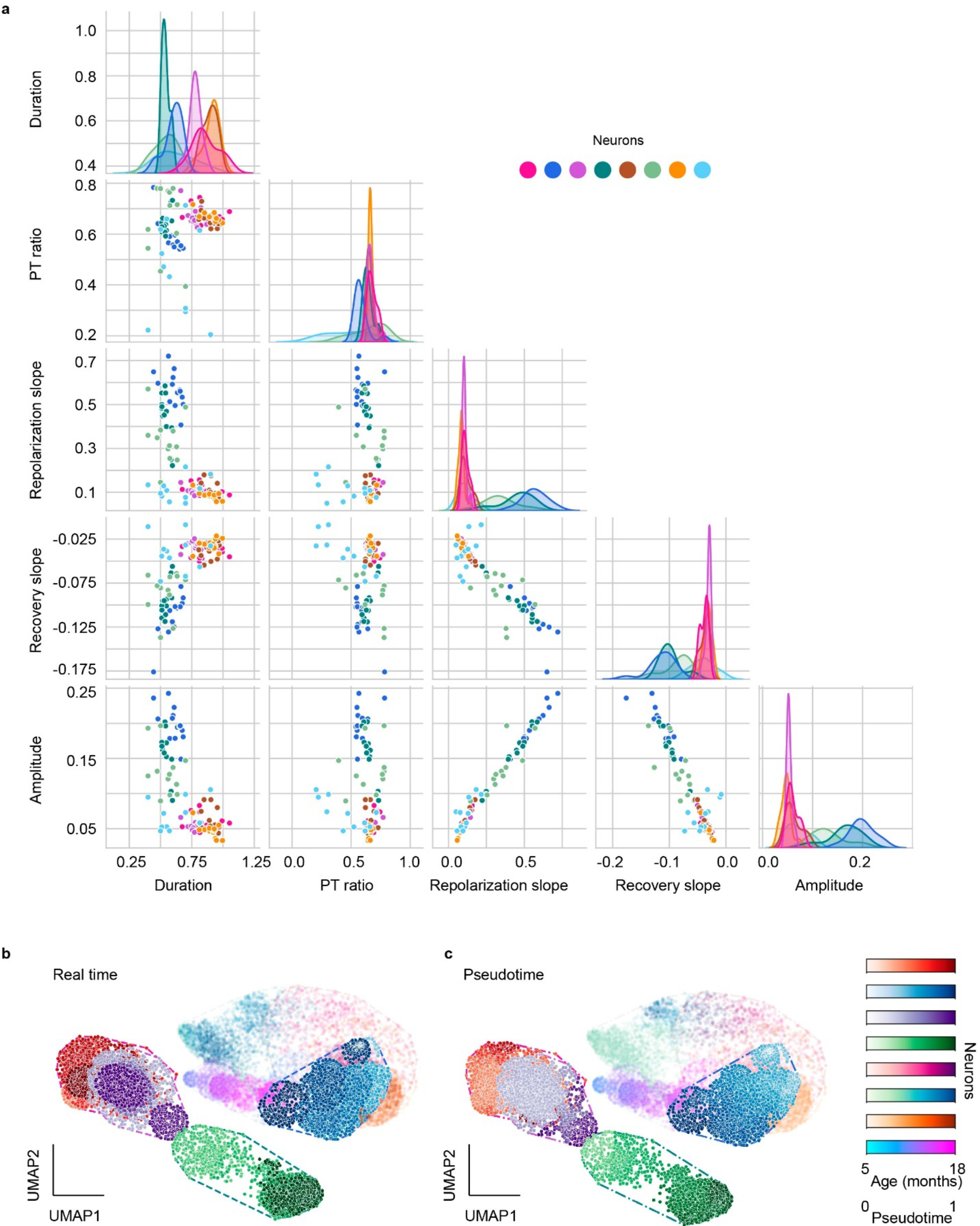
Feature selection and pseudo time analysis for potential aging-associated analysis. **a**, Pairplot of 5 *AllenSDK* selected features shown in Fig. 4e-h. Dots represent mean values of paired features calculated over neuron clusters over the entire recording period. Diagonal subplots show neuron feature univariate distributions using kernel density estimators. **b**, **c**, Real time (**b**) and pseudotime (**c**) comparison of UMAP embedding time-evolution for neurons in Fig. 4. The x- and y axes denote the first and second UMAP dimensions, respectively. Each gradient color-coded cluster represents a distinct neuron in Fig. 4. The color bars show the mouse age time points from 5 to 18 months and the corresponding pseudo time from 0 to 1. Highlighted dots correspond to the representative neurons used in the aging analysis of Fig. 6. Delimiting lines are the convex hull of a neuron’s UMAP embedding time-evolution.

**Extended Data Fig. 10.**
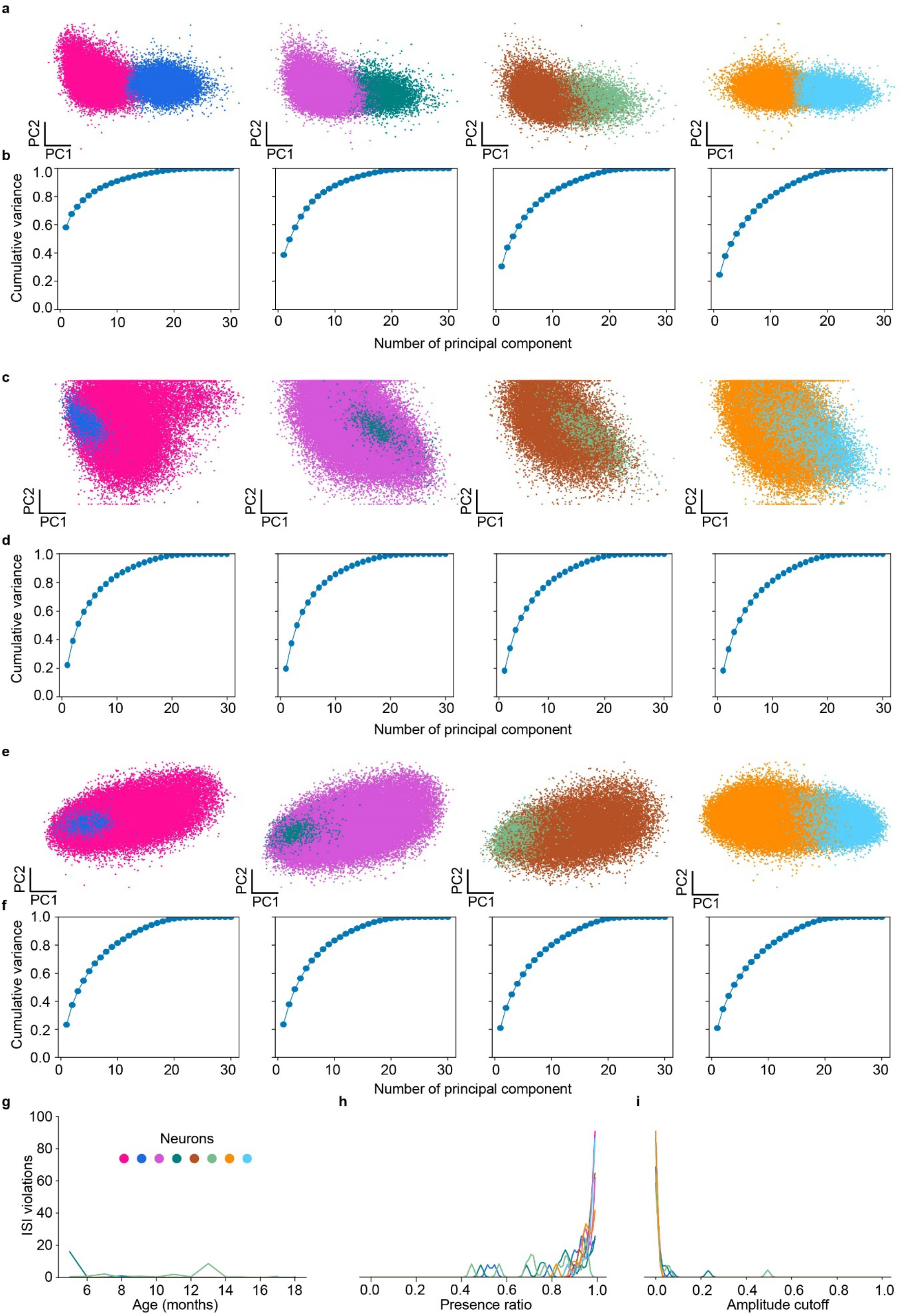
Quality assessment for PCA embeddings and neuron clusters. **a**, Principal Component (PC) embeddings of all recorded spikes without any normalization colored by neuron. **b**, Cumulative proportion of variance explained by top principal components of (**a**). **c**, **d**, Same as (**a**, **b**) with prior min-max normalization to scale spikes to (−1, 1) range. **e**, **f**, Same as (**a**, **b**) with prior standard scaling normalization per spike. Normalization decreases cluster separability in PC representation and lowers proportion of variance explained by the top two components by removing amplitude information from spikes. **g**, Interspike interval (ISI) violations in the absolute number of spikes calculated over all recordings for each recording month. **h**, Presence ratio smoothed density plot over all recording sessions per unit. **i**, Amplitude cutoff smoothed density plot over all recording sessions per unit.

## References

1 Gallego, J. A., Perich, M. G., Chowdhury, R. H., Solla, S. A. & Miller, L. E. Long-term stability of cortical population dynamics underlying consistent behavior. Nat. Neurosci. 23, 260–270 (2020).

2 Steinmetz, N. A. et al. Neuropixels 2.0: A miniaturized high-density probe for stable, long-term brain recordings. Science 372, eabf4588 (2021).

3 Dhawale, A. K. et al. Automated long-term recording and analysis of neural activity in behaving animals. eLife 6, e27702 (2017).

4 Schoonover, C. E., Ohashi, S. N., Axel, R. & Fink, A. J. P. Representational drift in primary olfactory cortex. Nature 594, 541–546 (2021).

5 Igarashi, K. M., Lu, L., Colgin, L. L., Moser, M.-B. & Moser, E. I. Coordination of entorhinal–hippocampal ensemble activity during associative learning. Nature 510, 143–147 (2014).

6 Dhawale, A. K., Wolff, S. B. E., Ko, R. & Ölveczky, B. P. The basal ganglia control the detailed kinematics of learned motor skills. Nat. Neurosci. 24, 1256–1269 (2021).

7 Wang, M. et al. Neuronal basis of age-related working memory decline. Nature 476, 210–213 (2011).

8 Grady, C. The cognitive neuroscience of ageing. Nat. Rev. Neurosci. 13, 491–505 (2012).

9 Meng, G. et al. High-throughput synapse-resolving two-photon fluorescence microendoscopy for deep-brain volumetric imaging in vivo. eLife 8, e40805 (2019).

10 Salatino, J. W., Ludwig, K. A., Kozai, T. D. Y. & Purcell, E. K. Glial responses to implanted electrodes in the brain. *Nat*. Biomed. Eng. 1, 862–877 (2017).

11 Ji, N. The Practical and fundamental limits of optical imaging in mammalian brains. Neuron 83, 1242–1245 (2014).

12 Yu, K. J. et al. Bioresorbable silicon electronics for transient spatiotemporal mapping of electrical activity from the cerebral cortex. Nat. Mater. 15, 782–791 (2016).

13 Chiang, C.-H. et al. Development of a neural interface for high-definition, long-term recording in rodents and nonhuman primates. Sci. Transl. Med. 12, eaay4682 (2020).

14 Song, E. et al. Flexible electronic/optoelectronic microsystems with scalable designs for chronic biointegration. Proc. Natl. Acad. Sci.116, 15398–15406 (2019).

15 Liu, Y. et al. Soft and elastic hydrogel-based microelectronics for localized low-voltage neuromodulation. Nat. Biomed. Eng. 3, 58–68 (2019).

16 Yin, R. et al. Soft transparent graphene contact lens electrodes for conformal full-cornea recording of electroretinogram. Nat. Commun. 9, 2334 (2018).

17 Zhang, J. et al. Stretchable transparent electrode arrays for simultaneous electrical and optical interrogation of neural circuits in vivo. Nano Lett. 18, 2903–2911 (2018).

18 Liu, J., et al. Syringe-injectable electronics. Nat. Nanotechnol. 10, 629–636 (2015).

19 Yang, X., et al. Bioinspired neuron-like electronics. Nat. Mater. 18, 510–517 (2019).

20 Fu, T.-M. et al. Stable long-term chronic brain mapping at the single-neuron level. Nat. Methods 13, 875–882 (2016).

21 Guan, S. et al. Elastocapillary self-assembled neurotassels for stable neural activity recordings. Sci. Adv. 5, eaav2842 (2019).

22 Kim, T.-i. et al. Injectable, cellular-scale optoelectronics with applications for wireless optogenetics. Science 340, 211–216 (2013).

23 Musk, E. An integrated brain-machine interface platform with thousands of channels. J. Med. Internet. Res.21,e16194 (2019)

24 He, F. et al. Multimodal mapping of neural activity and cerebral blood flow reveals long-lasting neurovascular dissociations after small-scale strokes. Sci. Adv. 6, eaba1933 (2020).

25 Sharp, A. A., Ortega, A. M., Restrepo, D., Curran-Everett, D. & Gall, K. In vivo penetration mechanics and mechanical properties of mouse brain tissue at micrometer scales. IEEE Trans. Biomed. Eng. 56, 45–53 (2008).

26 McInnes, L., Healy, J. & Melville, J. Umap: Uniform manifold approximation and projection for dimension reduction. Preprint at https://arxiv.org/abs/1802.03426 (2018).

27 Liu, W. et al. A survey of deep neural network architectures and their applications. Neurocomputing 234, 11–26 (2017).

28 Jeong, J.-W. et al. Wireless optofluidic systems for programmable in vivo pharmacology and optogenetics. Cell 162, 662–674 (2015).

29 Seo, K. J. et al. Transparent, flexible, penetrating microelectrode arrays with capabilities of single-unit electrophysiology. Adv. Biosyst. 3, 1800276 (2019).

30 Xie, C. et al. Three-dimensional macroporous nanoelectronic networks as minimally invasive brain probes. Nat. Mater. 14, 1286–1292 (2015).

31 Kozai, T. D. Y. et al. Ultrasmall implantable composite microelectrodes with bioactive surfaces for chronic neural interfaces. Nat. Mater. 11, 1065–1073 (2012).

32 Lu, L. et al. Soft and MRI compatible neural electrodes from carbon nanotube fibers. Nano Lett. 19, 1577–1586 (2019).

33 Guitchounts, G., Markowitz, J. E., Liberti, W. A. & Gardner, T. J. A carbon-fiber electrode array for long-term neural recording. J. Neural Eng. 10, 046016 (2013).

34 Rousche, P. J. et al. Flexible polyimide-based intracortical electrode arrays with bioactive capability. IEEE Trans. Biomed. Eng. 48, 361–371 (2001).

35 Minev, I. R. et al. Electronic dura mater for long-term multimodal neural interfaces. Science 347, 159–163 (2015).

36 Airaghi Leccardi, M. J. I., Vagni, P. & Ghezzi, D. Multilayer 3D electrodes for neural implants. J. Neural Eng. 16, 026013 (2019).

37 Barrese, J. C. et al. Failure mode analysis of silicon-based intracortical microelectrode arrays in non-human primates. J. Neural Eng. 10, 066014 (2013).

38 Chaure, F. J., Rey, H. G. & Quiroga, R. Q. A novel and fully automatic spike-sorting implementation with variable number of features. J. Neurophysiol. 120, 1859–1871 (2018).

39 Viswanathan, P. & Nieder, A. Visual receptive field heterogeneity and functional connectivity of adjacent neurons in primate frontoparietal association cortices. J. Neurosci. 37, 8919–8928 (2017).

40 Jia, X. et al. High-density extracellular probes reveal dendritic backpropagation and facilitate neuron classification. J. Neurophysiol. 121, 1831–1847 (2019).

41 Flurkey, K., M. Currer, J. & Harrison, D. E. The Mouse in Biomedical Research Ch.20 (eds James G. Fox et al.) 637–672 (Academic Press, Burlington, 2007).

42 Zhao, S. et al. Full activation pattern mapping by simultaneous deep brain stimulation and fMRI with graphene fiber electrodes. Nat. Commun. 11, 1788 (2020).

43 Schmitzer-Torbert, N., Jackson, J., Henze, D., Harris, K. & Redish, A. D. Quantitative measures of cluster quality for use in extracellular recordings. Neuroscience 131, 1–11 (2005).

44 Rousseeuw, P. J. Silhouettes: A graphical aid to the interpretation and validation of cluster analysis. J. Comput. Appl. Math. 20, 53–65 (1987).

45 Wang, Q., Webber, R. M. & Stanley, G. B. Thalamic synchrony and the adaptive gating of information flow to cortex. Nat. Neurosci. 13, 1534–1541 (2010).

46 Breiman, L. Random Forests. Mach. Learn. 45, 5–32 (2001).

47 Cao, J. et al. The single-cell transcriptional landscape of mammalian organogenesis. Nature 566, 496–502 (2019).

48 Zhao, S. et al. Graphene encapsulated copper microwires as highly MRI compatible neural electrodes. Nano Lett. 16, 7731–7738 (2016).

49 Traag, V. A., Waltman, L. & van Eck, N. J. From Louvain to Leiden: guaranteeing well-connected communities. Sci. Rep. 9, 5233 (2019).

